# Early Stem Cell Aging in the Mature Brain

**DOI:** 10.1101/654608

**Authors:** Albina Ibrayeva, Maxwell Bay, Elbert Pu, David Jörg, Lei Peng, Heechul Jun, Naibo Zhang, Daniel Aaron, Congrui Lin, Galen Resler, Axel Hidalgo, Mi-Hyeon Jang, Benjamin D. Simons, Michael A. Bonaguidi

## Abstract

Stem cell dysfunction drives many age-related disorders. Identifying mechanisms that initially compromise stem cell function represent early targets to enhance stem cell behavior later in life. Here, we pinpoint multiple factors that disrupt neural stem cells (NSC) in the adult hippocampus. We find that NSCs exhibit asynchronous maintenance by identifying short-term (ST-NSC) and intermediate-term NSCs (IT-NSCs). ST-NSC divide rapidly to generate neurons and deplete in the young brain. Meanwhile, multipotent IT-NSCs persist for months, but are pushed out of homeostasis by lengthening quiescence. Single cell transcriptome analysis of deep NSC quiescence revealed several hallmarks of biological aging in the mature brain and identified tyrosine-protein kinase Abl1 as an NSC pro-aging factor. Treatment with the Abl-inhibitor Imatinib increased NSC proliferation without impairing NSC maintenance in the middle-aged brain. Further intersectional analysis of mature NSC with old epidermal, hematopoietic and muscle stem cell transcriptomes identified consensus changes in stem cell aging. Our study elucidates multiple origins of adult neurogenesis decline and reveals that hippocampal NSCs are particularly vulnerable to a shared stem cell aging signature.

## INTRODUCTION

Aging is the progressive loss of physiological function due to accumulating cellular damage (Kennedy et al., 2014; López-Otín et al., 2013). This process occurs at a gradual and asynchronous rate where specific cells and then organ systems lose homeostasis (Almanzar et al., 2020; Negredo et al., 2020; Schaum et al., 2020). Aging has been historically investigated in chronological terms, with the majority of studies comparing young and old organisms. More recently, approaches to prevent aging at younger ages have become more common (Belsky et al., 2015; Gillman, 2005). Mechanisms that initiate aging are thought to represent early targets for promoting tissue resiliency throughout life. However, progress in slowing declining tissue function has been hampered by an inability to determine when and which cells begin to exhibit biological aging (López-Otín et al., 2013; Solanas et al., 2017).

Stem cells represent a key pillar of aging (Kennedy et al., 2014). Stem cells are present in many adult tissues and are responsible for tissue generation throughout life. In order to do so, they must balance new cell production with stem cell maintenance. However, this equilibrium is progressively lost during chronological aging in several tissues including the brain, blood, muscle and skin (Clevers, 2015; Goodell and Rando, 2015; Liu et al., 2013; Ziebell et al., 2018). Loss of stem cell function contributes to degeneration in tissue integrity and a reduced capacity for regeneration upon injury (Sahin and DePinho, 2010). Several lines of evidence indicate that stem cells exhibit molecular aging. Stem cells from older organisms display somatic mutations, epigenetic erosion, impaired cellular metabolism, oxidative stress and proteostasis damage (Berman et al., 2017; Goodell and Rando, 2015; Limke et al., 2002; López-Otín et al., 2013; Mattson et al., 1999; Southall et al., 2013). Yet, it remains unclear which stem cell aging molecular signatures are broadly shared among multiple tissues (Keyes and Fuchs, 2018; Sato et al., 2017; Solanas et al., 2017).

Neural stem cells represent a cell type that could be particularly vulnerable to cellular aging. Adult neurogenesis persists throughout life in the subgranular zone (SGZ) in the dentate gyrus of the hippocampus and in the subventricular zone (SVZ) along the lateral ventricles (Kirschen et al., 2019; Kuhn et al., 2018; Obernier and Alvarez-Buylla, 2019; Toda and Gage, 2018). In the adult hippocampus, neural stem cells (NSCs) give rise to new dentate granule neurons and astrocytes through a sequential process of cell cycle entry and generation of intermediate progenitor cells (Bonaguidi et al., 2011; Bond et al., 2015). New neurons play critical roles in regulating neural plasticity as well as cognitive and affective behaviors, whereas deficits in adult hippocampal neurogenesis have been implicated in age-related brain disorders (Anacker and Hen, 2017; Choi et al., 2018; Peng and Bonaguidi, 2018). While hippocampal neurogenesis persists throughout adulthood, this process is highly compromised during chronological aging. Aged animals have significantly lower neural stem cell numbers, stem cell proliferation, neuronal differentiation and newborn neuron survival compared to younger animals (Encinas et al., 2011; Heine et al., 2004; Kuhn et al., 1996; Ziebell et al., 2018). Further, NSCs in old animals exhibit molecular hallmarks of cellular aging including deficits in proteostasis and receive high levels of inflammation (Harris et al., 2020; Kalamakis et al., 2019; Leeman et al., 2018; Negredo et al., 2020). Yet, the hippocampus experiences a loss of neurogenesis early in the mature brain of rodents (Ben Abdallah et al., 2010; Morgenstern et al., 2008) and by middle-age in humans (Knoth et al., 2010; Moreno-Jiménez et al., 2019; Spalding et al., 2013). This decline is accompanied by epigenetic loss of DNA demethylation (Gontier et al., 2018), suggesting NSCs could molecularly age at early stages of chronological aging.

However, cellular origins driving early neurogenesis decline remain unclear. In one scenario, NSCs have been found to drop in number in the young hippocampus (Gontier et al., 2018; Lugert et al., 2012). This is supported by the presence of a finite NSC pool containing limited self-renewal capability (Encinas et al., 2011; Pilz et al., 2018). Consistently, forced accelerated neurogenesis (Ehm et al., 2010; Jones et al., 2015; Renault et al., 2009) and pathological conditions (Mu and Gage, 2011; Sierra et al., 2015) can result in premature NSC depletion. In another scenario, studies (Bonaguidi et al., 2011; Dranovsky et al., 2011; Licht et al., 2016) have shown that self-renewing NSCs are maintained for prolonged periods and neurogenesis instead could decline during aging due to an increase in NSC quiescence (Hattiangady and Shetty, 2008; Ziebell et al., 2018). While NSCs as a population clearly undergo early age-related changes, when and how specific NSC subpopulation begin to exhibit dysfunction remain unclear. In this study, we used single cell approaches to investigate the cellular and molecular mechanisms that initially compromise adult neurogenesis. We determined that NSC subpopulations undergo asynchronous decline and exhibit molecular aging in the mature hippocampus. We further identify a stem cell aging signature shared across multiple aged tissues. Finally, we show that targeting molecular aging in the middle-aged hippocampus can partially overcome age-related NSC dysfunction.

## RESULTS

### Asynchronous NSC decline during aging

We first investigated how adult NSC numbers change over time at the population level. The dentate gyrus (DG) of the hippocampus was harvested from wild-type mice and processed by immunofluorescence to reveal NSC behavior across chronological age. Nestin staining of radial-glial like neural stem cells indicated that NSC number decreases over time (Figure 1A). Consistent with prior observations, NSC loss is most pronounced during the transition from the young to mature and middle-aged brain (Gontier et al., 2018). We further processed neural stem cells for the cell cycle marker MCM2 to quantify age-related changes in radial glia like NSC proliferation. Similar to findings in older mice (Encinas et al., 2011; Hattiangady and Shetty, 2008; Ziebell et al., 2018), remaining NSCs become more quiescent early in the mature hippocampus (Figure 1C, S1A). Therefore, radial-glia like NSC number declines over time and remaining NSCs do not divide as frequently at the population level (Figure 1A-C, S1A).

**Figure 1.**
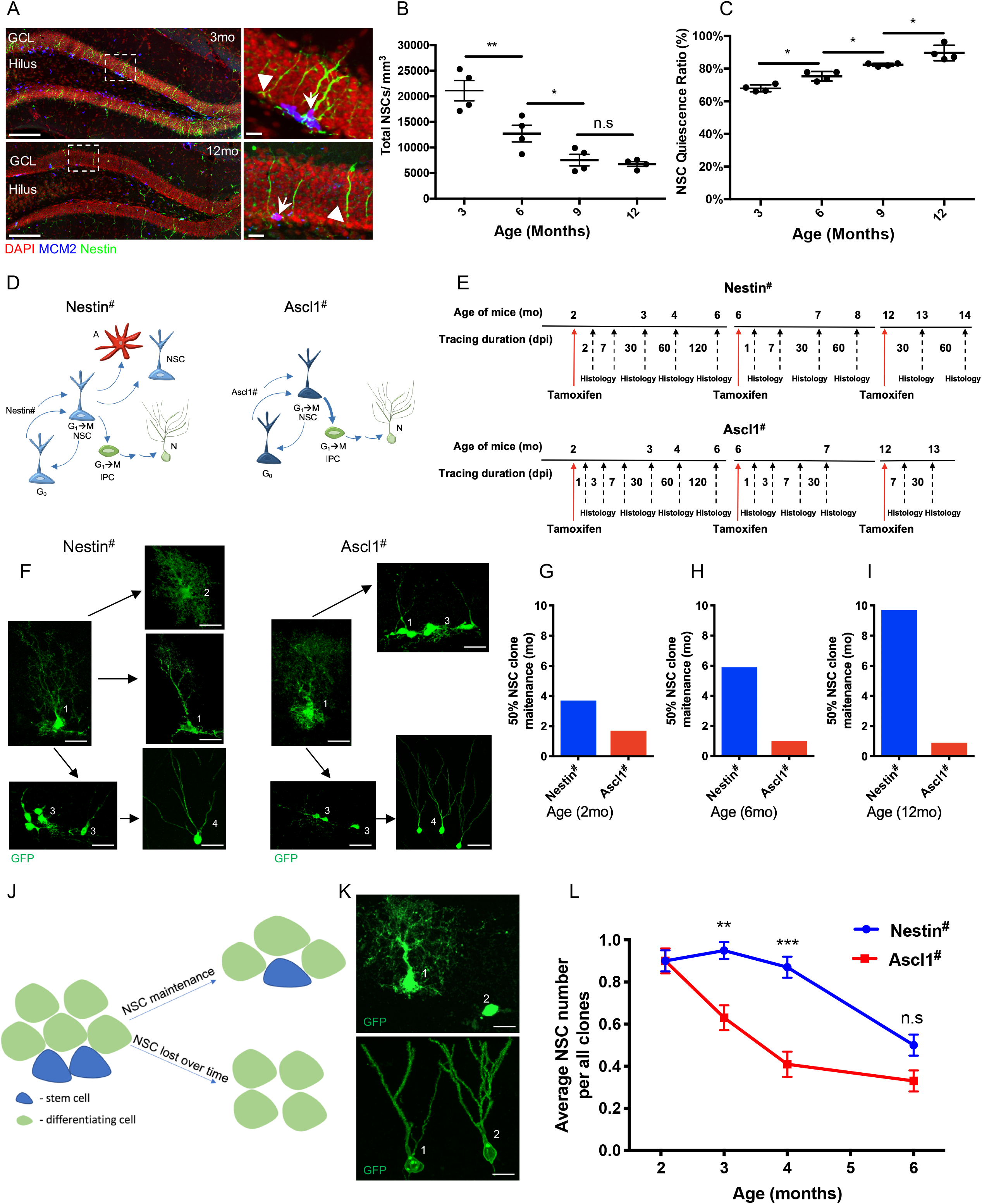
Asynchronous neural stem cell (NSC) decline during aging. (A) Sample confocal images showing immunofluorescence for NSCs (Nestin^+^) and cell proliferation (Mcm2^+^) in the dentate gyrus of 3-month-old and 12-month-old mouse hippocampus. Arrow - Nestin^+^Mcm2^+^ aNSC, Arrowhead - Nestin^+^Mcm2^+^ qNSC. Scale bar, 100μm (25μm for the insert). (B) Quantification of the total NSC number in the dentate gyrus across ages. N=4-6 mice; Values represent mean ± SEM. *p<0.05, **p<0.01, ***p<0.001, n.s - not significant; ANOVA with Bonferroni post-hoc test. (C) Quantification of the percentage of quiescent NSCs (Nestin^+^ MCM2^−^) among total NSCs in the dentate gyrus across ages. N=4-6 mice. Values represent mean ± SEM. *p<0.05, **p<0.01, ***p<0.001, n.s - not significant; ANOVA with Bonferroni post-hoc test. (D) Cartoon of NSC subpopulations in the adult hippocampus. (left) Nestin::CreER (Nestin#)-labeled multipotential NSCs. (right) Developmental-like NSCs labeled by Ascl1::CreER (Ascl1#). A=astroglial lineage, IPC=intermediate progenitor cell, N=neuronal lineage, G_0_=quiescent state, G_1_->M=active state. (E) Experimental design of *in vivo* single cell clonal lineage tracing for Nestin#-NSC and Ascl1#-NSC subpopulations, results in Table S1 and S4. (F) Sample confocal images of NSC subpopulations in the adult hippocampus. (left) Nestin::CreER (Nestin#)-labeled multipotential NSCs. (right) Developmental-like NSCs labeled by Ascl1::CreER (Ascl1#). NSC = neural stem cell (1), A= astroglial lineage (2), intermediate progenitor cells (IPCs) (3) and neurons (4). Scale bar, 10μm (G) Nestin#-NSCs display greater 50% NSC clone maintenance (the time until 50% of clones exhibit NSC depletion) than Ascl1#-NSCs in 2 month-old mice. N=29-104 clones, see details in Table S4; (H) Nestin#-NSCs display greater 50% NSC clone maintenance (the time until 50% of clones exhibit NSC depletion) than Ascl1#-NSCs in 6 month-old mice. N=29-104 clones, see details in Table S4; (I) Nestin#-NSCs display greater 50% clonal maintenance (the time until 50% of clones exhibit NSC depletion) than Ascl1#-NSCs in 12 month-old mice. N=29-104 clones, see details in Table S4; (J) Cartoon depicting clonal maintenance and depletion. (K) Sample confocal images of a maintained clone containing NSC (1) and intermediate progenitor cells (IPCs) (2); and a depleted clone containing neurons (1) and (2). Scale bar, 10μm (L) Quantification of NSC homeostasis duration. Ascl1#-NSCs are rapidly depleted, while Nestin#-NSCs maintain as stem cells for months before eventually differentiating. Values represent mean ± SEM. (N = 29-104 clones, see details in Table S4). Values represent mean ± SEM. *p<0.05, **p<0.01, ***p<0.001, n.s - not significant; two-way ANOVA with Tukey’s multiple comparisons test.

Various observations of NSC behavior have been made in the adult hippocampus. Some NSCs are short-lived and divide rapidly to mainly produce granule neurons – exhibiting “division-coupled differentiation” behavior (Encinas et al., 2011) or “developmental-like model” behavior (Pilz et al., 2018). Meanwhile, some studies (Bonaguidi et al., 2011; Dranovsky et al., 2011; Jang et al., 2013; Licht et al., 2016) have advanced a stem cell maintenance model where NSCs are longer lived, slowly cycle in and out of quiescence, can generate neurons, astrocyte and self-renew. We reasoned the existence of NSC subpopulations within the radial glial-like NSC pool masks the ability to precisely uncover which mechanisms contribute to NSC dysfunction over time (Bonaguidi et al., 2012, 2016). We therefore performed *in vivo* single cell clonal lineage tracing and computational modeling of *Nestin-CreER^T2^* (Nestin#) and *Ascl1-CreER^T2^* (Ascl1#) mice during physiological conditions at 2-, 6- and 12 months of age (young, mature and middle-age, respectively) according to our prior approaches (Bonaguidi et al., 2011) (Figure 1D-F, S1E-G). We found that clonally labelled Ascl1#-NSCs behave as a short-term neuronal fate-biased subpopulation, consistent with a developmental-like program (Pilz et al., 2018) (Figure 1D-I, S1C-D, S2D-E). Analysis of Ascl1#-NSC clonal number over time demonstrates consistent short-term stem cell maintenance in the young, mature and middle-aged hippocampus (Figure 1J-L, S1B). Meanwhile, individual NSCs marked by Nestin^#^-NSCs are longer-lived, slow cycling, generate neurons and astrocytes, and self-renew (Figure 1D-I, S1C-D, S2A-C). Importantly, these Nestin#-NSCs are homeostatic in the young brain for a few months – where the average number of NSC per clone is close to 1 - then transition out of homeostasis in roughly 4.5-months-old mice (Figure 1J-L). Computational modeling revealed that Nestin#-NSC clones are better maintained than Ascl1#-NSCs clones in the young, mature and middle-aged hippocampus. Further, remaining Nestin#-NSCs display progressively increased NSC maintenance at older ages compared to younger ages (Figure 1G-I, S1C-D). These data show consistent short-lived and longer-lived properties among NSC subpopulations across multiple ages. Thus, Nestin#-NSCs represent an intermediate-term NSC subpopulation (IT-NSCs) and can serve as a platform to pinpoint mechanisms that mediate the initial loss of stem cell homeostasis in the mature brain.

### Increased quiescence drives NSC loss of homeostasis

To begin exploring why Nestin#-NSCs lose homeostasis, we considered cellular mechanisms that would lead to declining NSC numbers. NSCs are homeostatic when their rate of symmetric self-renewal (expansion) balances their rate of differentiation (Figure 2A). We reasoned that NSCs either increase their differentiation or slow their expansion during their transition out of homeostasis. We first tested NSC clonal depletion in 2 to 6-months-old mice to analyze the rate of NSC differentiation. Clone maintenance was calculated by identifying the percentage of clones per time point that contain 1 + number of NSCs. As expected (Pilz et al., 2018), Ascl1#-NSCs displayed a rapid initial depletion that slowed with time. Meanwhile, Nestin^#^-NSCs exhibited a constant and gradual loss of clones containing a single NSC, indicating that homeostasis decline does not occur due to accelerating NSC differentiation (Figure 2B). We then analyzed NSC-containing clones to identify age-related changes in the rate of NSC expansion. Ascl1^#^-NSCs displayed no appreciable expansion over time (Pilz et al., 2018). On the other hand, Nestin#-NSCs did expand in young mice according to predicted values that maintain homeostasis (Figure 2C). However, NSC expansion quickly began to diverge from levels needed to maintain homeostasis in approximately 4-month-old mice. Hence, Nestin#-NSCs lose homeostasis in the mature brain because of slowing NSC production while Ascl1#-NSCs do not exhibit homeostasis due to a lack of substantial NSC production in the young brain (Figure 2A-C).

**Figure 2.**
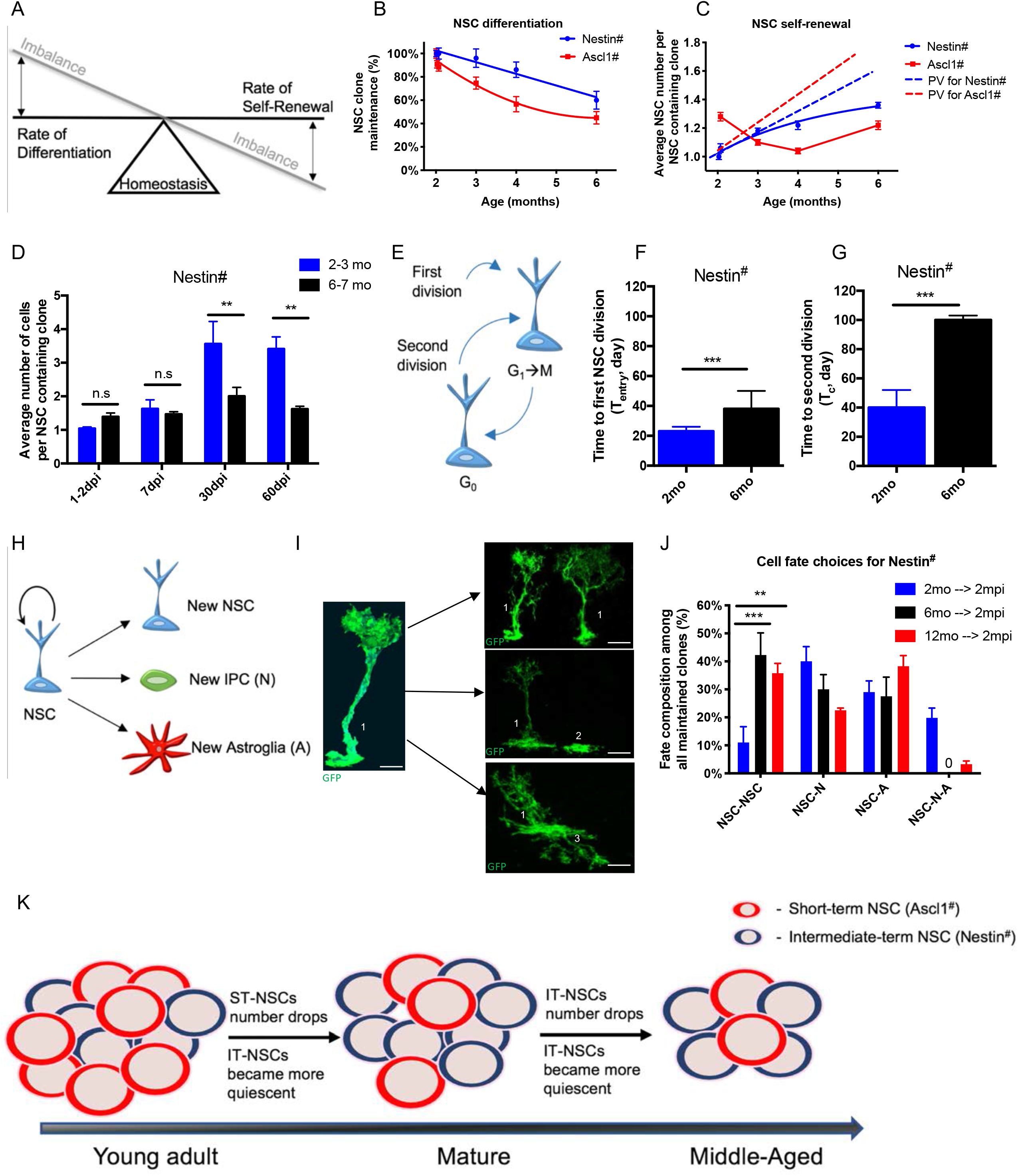
Increased quiescence drives NSC loss of homeostasis. (A) Cartoon illustrating NSC homeostasis and its imbalance by changes in NSC expansion or depletion. (B) Quantification of NSC clone maintenance in the mature hippocampus, Nestin#-NSC clones are depleted at a constant rate. (N = 30-69 clones, see details in Table S4). (C) Quantification of NSC self-renewal in the mature hippocampus, Nestin#-NSCs slow their expansion in approximately 4-month old mice indicating homeostatic imbalance. PV=predicted value to maintain NSC homeostasis. (N = 30-69 clones, see details Table S4). (D) Quantification of average number of cells per NSC-containing clones across 2- and 6-months old animals at multiple days post tamoxifen injection (detailed information Table S4). (N = 30-69 clones, see details Table S4; *p<0.05, **p<0.01, ***p<0.001, n.s - not significant; ANOVA with Bonferroni post-hoc test). (E) Cartoon of time to the first and second NSC divisions. (G_0_ = quiescence; G_1_ -> M = activation). (F) Quantification of Nestin#-NSC cell cycle entry based on lineage analysis from 2-month-old and 6-month-old mice. Values represent mean ± SEM. (N = 30-69 clones, see details Table S4; *p<0.05, **p<0.01, ***p<0.001, n.s - not significant; ANOVA with Bonferroni post-hoc test). (G) Quantification of Nestin#-NSC cell cycle re-entry based on lineage analysis from 2-month-old and 6-month-old mice. Values represent mean ± SEM (N = 30-69 clones, see details Table S4; *p<0.05, **p<0.01, ***p<0.001, n.s - not significant; ANOVA with Bonferroni post-hoc test). (H) Schematic illustration of NSC fate choices. NSC=neural stem cell; A=astroglial lineage; IPC=intermediate progenitor cell; N=neuronal lineage. (I) Representative confocal images of clonal NSC self-renewing fate choices. 1 – NSC; 2 – N (neuronal lineage); 3 – A (astroglial lineage). Scale bar, 10 μm. (J) Quantification of self-renewing cell fate choices outcome from Nestin#, clones acquired across multiple ages 2, 6- and 12-months old animals at 60 days (2 months) post tamoxifen injection (detailed information Table S1). NSC=neural stem cells; A=Astroglial lineage; N=neuronal lineage, 0 – no observed phenotype. Values represent mean ± SEM (*p<0.05, **p<0.01, ***p<0.001, n.s - not significant; ANOVA with Bonferroni post-hoc test); (K) Cartoon summary of the cellular mechanisms driving age-related NSC dysfunction.

NSC expansion occurs in a two-step process: transition from quiescence to cell cycle and then symmetric cell division. To determine which stem cell behavior drives the loss of NSC homeostasis, we performed clonal expansion, cell cycle kinetics and cell fate analyses on Nestin#-NSC clones. Two-months-old, 6-months-old and 12-months-old Nestin^#^ mice were traced up to 4 months, 2 months, and 2 months past tamoxifen injection, respectively (Table S4). We first quantified the average number of cells per NSC-containing clone to assess clonal expansion. NSC clone sizes at older ages were smaller compared to younger ages. Interestingly, the initial decline in clonal expansion occurred at 6 months of age and was consistently lower at 12 months of age (Figure 2D, S1I-J). We then assessed NSC quiescence by calculating the time to cell cycle entry and re-entry according to power-law decay fitting of clonal tracings (Figure 2E, (Pilz et al., 2018)). Both durations to the first and second division increased in Nestin#-NSCs between 2- and 6-month old mice (Figure 2F-G). These findings suggest that Nestin#-NSCs display a substantial increase in quiescence in the young hippocampus and enter a “deeper quiescent” state in the mature brain (van Velthoven and Rando, 2019). To determine age-related changes in cell fate choices, we followed the generation of each lineage type (neuronal, astroglial, NSCs) and lineage cell type (neuronal intermediate progenitor cells, immature neurons, mature neurons; astrocyte progenitor cells, astrocytes) within the Nestin^#^-NSC clones. Fate choices were calculated for each cell division among the neuronal, astroglial and NSC lineages (Figure 2H-J, S1H). As previously reported (Bonaguidi et al., 2011), Nestin#-NSCs predominately make neuronal and astroglial asymmetric fate choices in the young 2-month-old hippocampus. However, we found that symmetric cell divisions predominantly occur in the 6-month-old mature brain at the expense of neuronal asymmetric divisions (Figure S1H). Intriguingly, the age-related changes in Nestin-NSCs cell fate choices that occur at 6 months of age are preserved in middle-aged 12 months old mice (Figure 2J, S1H). Together, these observations indicate that Nestin#-NSCs prolong their quiescence with each division, which drives them from homeostasis in the mature hippocampus. Further, they then switch to symmetric cell fate choice after NSC homeostasis has been lost in the mature brain (Figure 2K).

### Quiescent NSC molecular aging in the mature hippocampus

NSCs are a rare cell population in the adult hippocampus that transition between quiescent and active states (Bonaguidi et al., 2011). While molecular signatures distinguishing quiescent from active NSCs have been well documented in the SVZ and hippocampus (Codega et al., 2015; Dulken et al., 2017; Llorens-Bobadilla et al., 2015; Shin et al., 2015), age-related changes within NSC quiescence have been more difficult to identify (Kalamakis et al., 2019; Leeman et al., 2018). Nestin#-NSCs are homeostatic in the young hippocampus for a few months, then exit homeostasis at around 4.5-months-old mice due to an increase in quiescence (Figures 1–2). To uncover molecular mechanisms of deepening NSC quiescence, we performed single cell RNA sequencing (scRNA-seq) (Figure 3A, S3A). NSCs and their immediate progeny were isolated using 2 months-old and 4.5 months-old Nestin::CFP mice (Encinas et al., 2006), and immediately enriched using FACs (Figure S3B). Nestin::CFP captures most NSCs, including shortterm and intermediate-term subpopulations.

**Figure 3.**
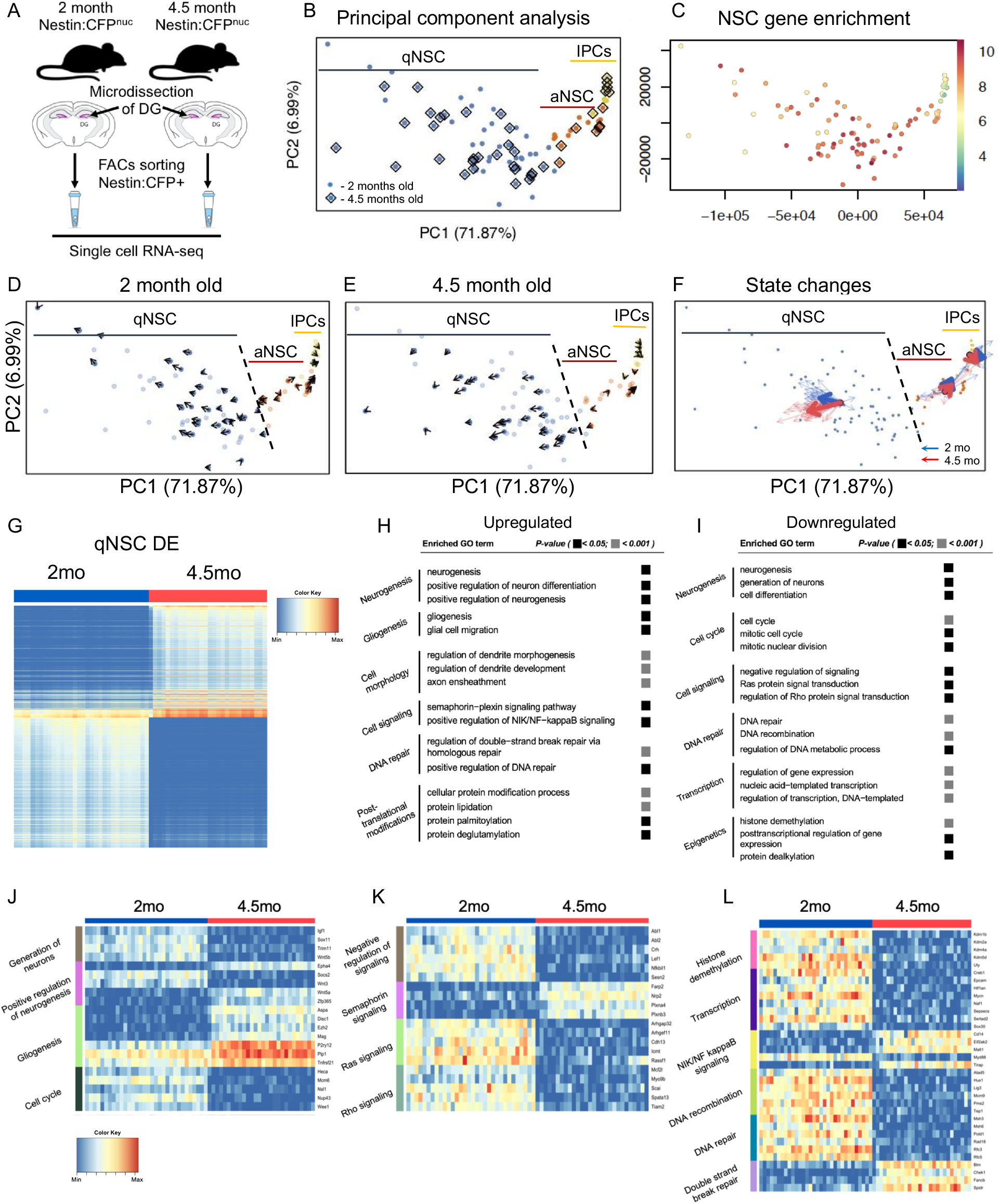
Quiescent NSC aging in the mature hippocampus. (A) Schematic illustration of the single cell RNA sequencing approach. (B) Principle component (PC) analysis of NSC transcriptomes from 2- and 4.5-month-old mice. (C) NSCs identified by gene enrichment analysis using Nestin^+^, Vimentin^+^, Fabp7^+^, Aldoc^+^, Apoe^+^ markers. (D) RNA velocity defines single cell future states (arrows) identified in quiescent NSCs (qNSC), active NSCs (aNSCs) and IPCs from 2-month-old hippocampus. (E) RNA velocity defines single cell future states (arrows) identified in qNSCs, active aNSCs and IPCs from 4.5-month-old hippocampus. (F) RNA velocity analysis reveals NSCs more resistant to activate in 4.5-month-old mice, whereas some qNSCs from 2-month-old mice will enter cell cycle. (G) Quiescent NSC differential expression (DE) heat map between 2- and 4.5-months old mice. (H) Top upregulated gene ontology (GO) terms between quiescent NSCs derived from 2-month-old and 4.5-months-old mice identify molecular aging. FDR-corrected p-values are shown. (I) Top downregulated gene ontology (GO) terms between quiescent NSCS derived from 2-month-old and 4.5-months-old mice identify molecular aging. FDR-corrected p-values are shown. (J) Heat map of Neurogenesis, Gliogenesis and Cell cycle transcript changes between quiescent NSCs derived from 2-month-old and 4.5-months-old mice. (K) Heat map of Semaphorin, Ras and Rho signaling and signaling transcript changes between quiescent NSCs derived from 2-month-old and 4.5-months-old mice. (L) Heat map of Histone Demethylation, Transcription, NIK/NF kappaB signaling, DNA recombination, DNA repair and Double strand break repair transcript changes between quiescent NSCs derived from 2-month-old and 4.5-months-old mice.

Single-cell cDNA libraries were built using a miniaturized SMART-Seq4 protocol, barcoded, multiplexed and sequenced using paired-end NextSeq 550 (Figure 3A, S3). Sequencing reads were aligned using STAR (Dobin et al., 2013) and expression estimates were computed from aligned reads with RSEM (Li and Dewey, 2011). After gene filtering, 14,750 genes were detected across cells, where the average cell expressed 3,600 detected genes to permit a deep exploration of NSC transcriptional dynamics within quiescence (Table S5). We distinguished NSCs (Nestin, Vimentin, Apoe, Fabp7) from intermediate progenitor cells (IPCs: Eomes/Tbr2, Sox11, Stmn1) and identified NSCs in quiescent (Aldoc, Aqpr4, Id3, Hes1) or active states (PCNA, Mcm7, Cdk4, Cdk6) using well established markers (Shin et al., 2015; Urbán et al., 2019) (Figure 3C, S4A-C). Importantly, both Ward minimum variance hierarchical clustering and principal component analysis (PCA) showed shared cell-state-specific expression profiles between the two time points, indicating that global transcriptomic differences are greater between quiescent and active NSC states than within NSC states cells from 2-month-old and 4.5-month-old mice (Figure 3B, S4D-F).

Next, we probed for changes within quiescence between 2-month-old and 4.5-month-old NSCs using RNA velocity. RNA velocity takes separate observations of unspliced and spliced reads to predict the future gene expression state of individual cells and reveal lineage progression (La Manno et al., 2018). RNA vectors in our analysis indicated that most NSCs remain quiescent but when activated NSCs either differentiate or return to quiescence (Figure 3D-F). We did not detect age-related differences in activated NSCs or IPCs isolated from 2-month-old and 4.5-month-old mice. Interestingly, we found that NSCs in quiescence are more likely to move away from activation in 4.5-month-old mice compared to 2-month-old mice (Figure 3F), again implicating that NSCs begin to enter deeper quiescence in the mature hippocampus.

We then sought to identify transcriptomic changes associated with this NSC deep quiescence. 493 upregulated and 576 downregulated genes were differentially expressed (DE) in quiescent NSCs from 4.5-month-old mice compared to 2-month-old mice (Figure 3G, S4D-F, Table S8). We then used TopGo (Alexa et al., 2006) to represent differentially expressed transcripts (DEs) by their associated biological processes. Consistent with our lineage tracing experiments (Figure 2), gene ontology revealed changes in neurogenesis (*Socs2, Sox11, Igf1, Wnt3, Epha4*), a gain in self-renewal/gliogenesis (*Ezh2, Disc1, Mag, Plp1*) and a loss of cell cycle (*Wee1, Nsl1, Mcm6, Heca*). In addition, we observed an overall increase in cell signaling (Abl1, Abl2, Crh, Lef1) and alterations in processes known to regulate adult NSC quiescence or progenitor proliferation including semaphorin signaling (*Plxna4, Plxnb3, Nrp2, Farp2*), Ras signaling (*Arhger11, Arhgep32, Rassf1, Cdh13, Icmt*) and Rho signaling (*Spata13, Myo9b, Tiam2, Mcf2l, Scai*) (Chavali et al., 2018; Jongbloets et al., 2017; Li et al., 2012) (Figure 3J-L, S4G-H, Table S8). Strikingly, the remainder of detected terms are molecular hallmarks of an intertwined process that drives cellular aging (Kennedy et al., 2014; López-Otín et al., 2013). These factors include epigenetic dysregulation from histone demethylation (*Kdm1b, Kdm2a, Kdm4a, Kdm5d, Uty*), downregulation of transcription (*Sox30, Mycn, Creb1, Hif1an, Sertad2, Epcam*), inflammation due to NIK/NF-kappaB signaling (*Cd14, Malt1, Eif2ak2, Tirap, Myd88*) and cellular stress from loss of DNA recombination (*Atad5, Hus1, Pms2, Lig3, Mcm9*), DNA repair (*Sesn2, Msh3, Msh6, Rfc3, Rfc5, Rad18*) and increased double-strand break repair (*Chek1, Blm, Spidr, Fancb*) (Figure 3H-L, S4G-H, Table S8). Therefore, functional changes occurring in deep NSC quiescence are associated with early molecular aging in the mature hippocampus.

### Consensus signatures of stem cell aging

Many aged tissues display functional decline due, in part, to stem cell dysfunction. Aging can compromise general stem cell fitness: their capacity to self-renew, activate from quiescence, and produce specialized cell types (Goodell and Rando, 2015). However, it remains unclear which stem cell aging molecular mechanisms are shared among multiple tissues (Keyes and Fuchs, 2018; Sato et al., 2017; Solanas et al., 2017) Since NSCs from the mature hippocampus exhibit molecular hallmarks of aging (Figure 3), we compared NSC transcriptomes (merged quiescent and active states) with published advanced age (21-26 month-old) hematopoietic – hSC (Sun et al., 2014), muscle – mSC (Liu et al., 2013) and epidermal – eSC (Solanas et al., 2017) stem cell transcriptome datasets (Figure 4A, Table S9). Fisher’s exact tests were performed on respective DEs to determine the degree of age-associated gene expression convergence between NSCs and other stem cell compartments. DEs driving molecular aging in NSCs compared against DEs from chronological aging in hSC, mSC, eSC showed statistically significant gene-level convergence in both expression directions (Figure 4B, Table S9). Further pairwise analysis of gene ontology between NSCs from the mature brain and old stem cells from other tissues predicted functional convergence of aging (Figure 4C). The degree of overlap between NSCs and mSCs, NSCs and eSCs was greater at the functional (GO) level than gene level (Figure 4B-C) suggesting that hippocampal aging is driven by an expression program that, although involves specific genes which are not widely shared with other stem cell compartments, nonetheless converges on the same biological processes of aging.

**Figure 4.**
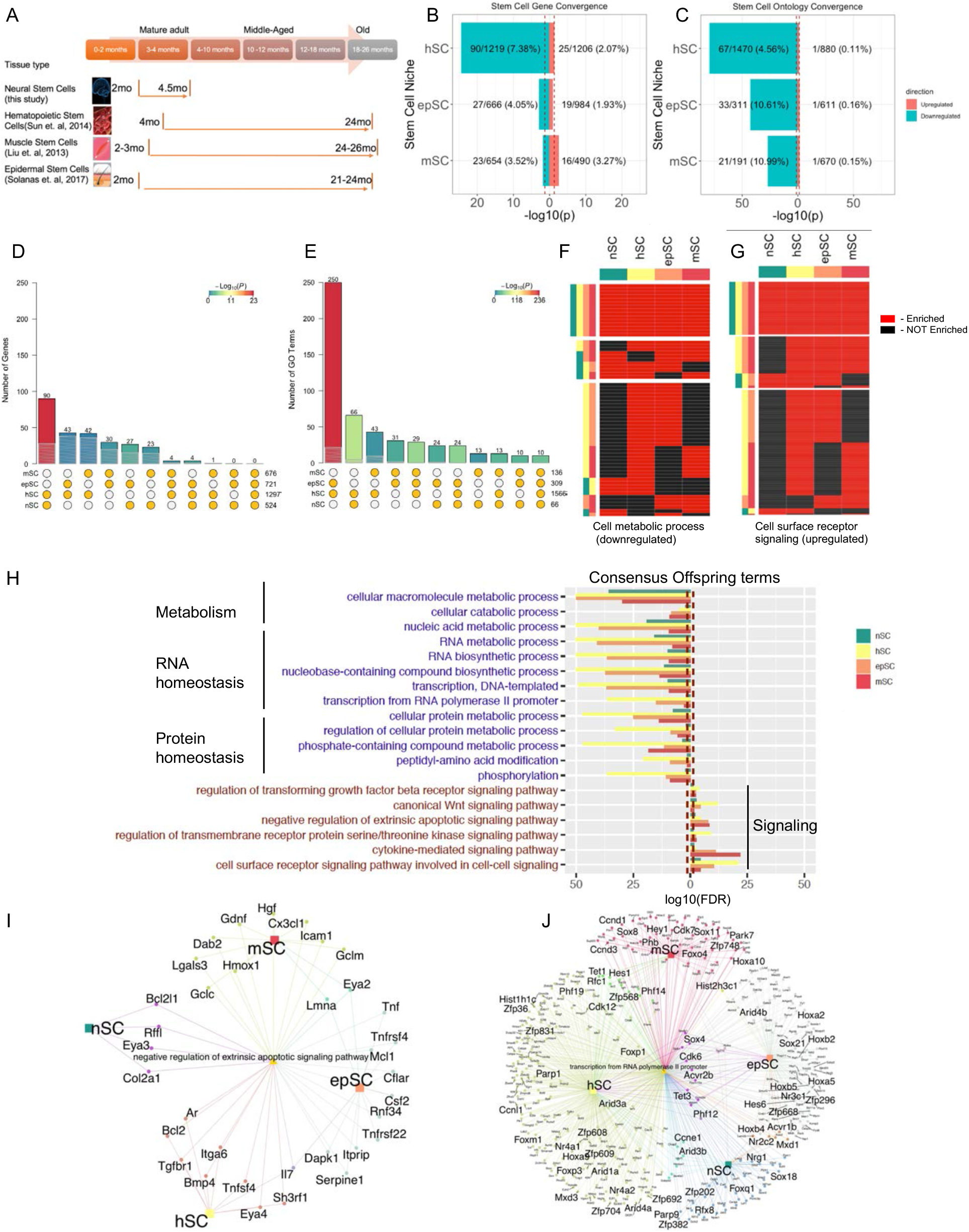
Consensus signatures of stem cell aging. (A) Experimental approach to compare stem cell aging using mature NSC transcriptomes compared to old hematopoietic (hSC), muscle (mSC) and epidermal (eSC) stem cell transcriptomes. (B) Pairwise stem cell gene convergence between NSCs and hSCs, mSCs, eSCs. FDR-corrected p-values are shown. (C) Pairwise stem cell ontology convergence between NSCs and hSCs, mSCs, eSCs. FDR-corrected p-values are shown. (D) Visualization of intersection among DE genes across multiple stem cell niches revealed by the SuperExactTest; p-value in descending order of statistical significance. (E) Visualization of intersection among GO terms across multiple stem cell niches revealed by the SuperExactTest; p-value in descending order of statistical significance. (F) Cell metabolic processes shared by various stem cell compartments. (G) Cell surface receptor signaling shared by various stem cell compartments. (H) Top consensus up- and downregulated offspring terms shared across four stem cell niches. Metabolism, RNA homeostasis and proteostasis are downregulated. Regulation of Wnt, TGFβ and apoptotic signaling is upregulated. FDR-corrected p-values are shown. (I) String network analysis showing unique or shared upregulated genes associated with negative regulation of apoptotic signaling pathway in NSCs, hSCs, mSCs, and eSCs. (J) String network analysis showing unique or shared downregulated genes associated with transcription from RNA polymerase II promoter in in NSCs, hSCs, mSCs, and eSCs.

We then investigated the presence of a consensus stem cell aging signature across 4 somatic stem cell compartments. Similar to prior reports (Keyes and Fuchs, 2018; Sato et al., 2017; Solanas et al., 2017), only a few genes (1-10) were shared among 3 stem cells and none were convergent among all 4 stem cells (Figure 4D). Indeed, permuted convergence of DE genes decreased monotonically when examining pairs, triads and quads of stem cell niches (Figure 4D). This observation suggests that stem cell aging acts on a set of genes that are largely unique to any respective niche. Interestingly, ontology-level convergence decreased between pairs and triads, but increased again when observing the convergence between all four stem cells (Figure 4E, Table S10-S11). This finding suggests molecular aging across stem cell niches recruit different but complementary genes which target common biological processes. In total, 18 gene ontology terms are differentially regulated by aging in all 4 stem cells (Figure S6C-D, Table S10-S11). These terms revealed general downregulation of metabolic processes and upregulation of cell signaling (Figure 4F-G). We then performed a more focused enrichment analysis on the DEs which were component genes of metabolism to understand which specific aspects were perturbed with aging. This analysis revealed a downregulation of genes associated with metabolic changes in RNA homeostasis including RNA biosynthesis, DNA-templated transcription and RNA polymerase II driven transcription; and metabolic changes in proteostasis such as control of peptidyl-amino acid modifications and phosphorylation (Figure 4F, H). Similarly, we performed a focused enrichment analysis on upregulated component genes of cell signaling. We identified changes in WNT signaling, TGFβ signaling and cell death signaling pathways (Figure 4G, H, Table S12). Thus, our results suggest that stem cell aging is driven by a general loss of intrinsic factors and a gain in regulation of extrinsic factors.

Finally, we generated string networks to visualize the gene expression changes within biological processes that are associated with molecular aging across the 4 stem cell data sets (Figure 4I-J). Different members of shared gene families were involved in aging across stem cells in every investigated ontology term among cell signaling, RNA homeostasis and proteostasis (Figures S5–7). Specifically, Eya transcription factor paralogs were downregulated in NSCs (Eya3), hSCs (Eya 4), mSCs and eSCs (Eya 2) (Figure 4I). In addition, regulators of RNA polymerase II were downregulated in all 4 stem cells, including members of *Tet, Fox, Sox* and *Hox* families (Figure 4J, Table S12). Thus, we have identified gene families that play complementary roles in stem cell aging across 4 tissues. Further, changes in old hematopoietic, muscle and epidermal stem cells are present early in neural stem cells from the mature hippocampus.

### Imatinib partially restores NSC function in the middle-aged brain

Strategies to delay and potentially even reverse the aging process have recently been developed (Mahmoudi et al., 2019; Negredo et al., 2020). Since NSCs undergo early cellular and molecular aging in the mature brain (Figure 3–4), we hypothesized that targeting this process could overcome age-related NSC dysfunction later in life (Figure 2–3). Gene networks are powerful approaches with the ability to prioritize genes based upon their degree of connectedness to other genes and functions (Li and Horvath, 2007). We therefore probed for differentially expressed genes shared between 3 or more GO terms for target identification (Figure 5A). Our analysis identified *Abl1, Abl2, Igf1, Lef1, Per2* and *Nup62* as genes most connected to changes in age-related NSC function – including changes in signal transduction, gene expression, cell cycle, DNA repair and neurogenesis (Figure 5A). We decided to focus on targeting Abl1, because it was identified as one of the main hub nodes in age-related NSC network and has many context-dependent signaling functions, but an unknown role in NSC biology (Wang, 2014). Abl1 expression decreased between 2- and 4.5-months old mice at the transcriptome level under physiological conditions (Figure 5B). However, immunostaining for c-Abl (Abl1) within radial-glia like NSCs (c-Abl^+^Nestin^+^) revealed more prevalent c-Abl protein in NSCs with chronological age advancing from 2 months to 10 months-old (Figure 5C-D). We then sought to inhibit Abl with Imatinib as a strategy to target NSC cellular aging. Imatinib or vehicle control was infused by the hippocampal fimbria of 10-month-old mice for 6 days (Figure 3E). We found that Imatinib treatment decreases the prevalence of Abl1 protein in NSCs (Figure 5F-G). Interestingly, Nestin staining of radial-glial like NSCs with the cell cycle marker MCM2 revealed that Imatinib increases NSC proliferation in the middle-aged hippocampus to younger levels without altering NSC number (Figure 6A-D). Thus, targeting mechanisms associated with the initial loss of NSC homeostasis can overcome deep quiescence later in life.

**Figure 5.**
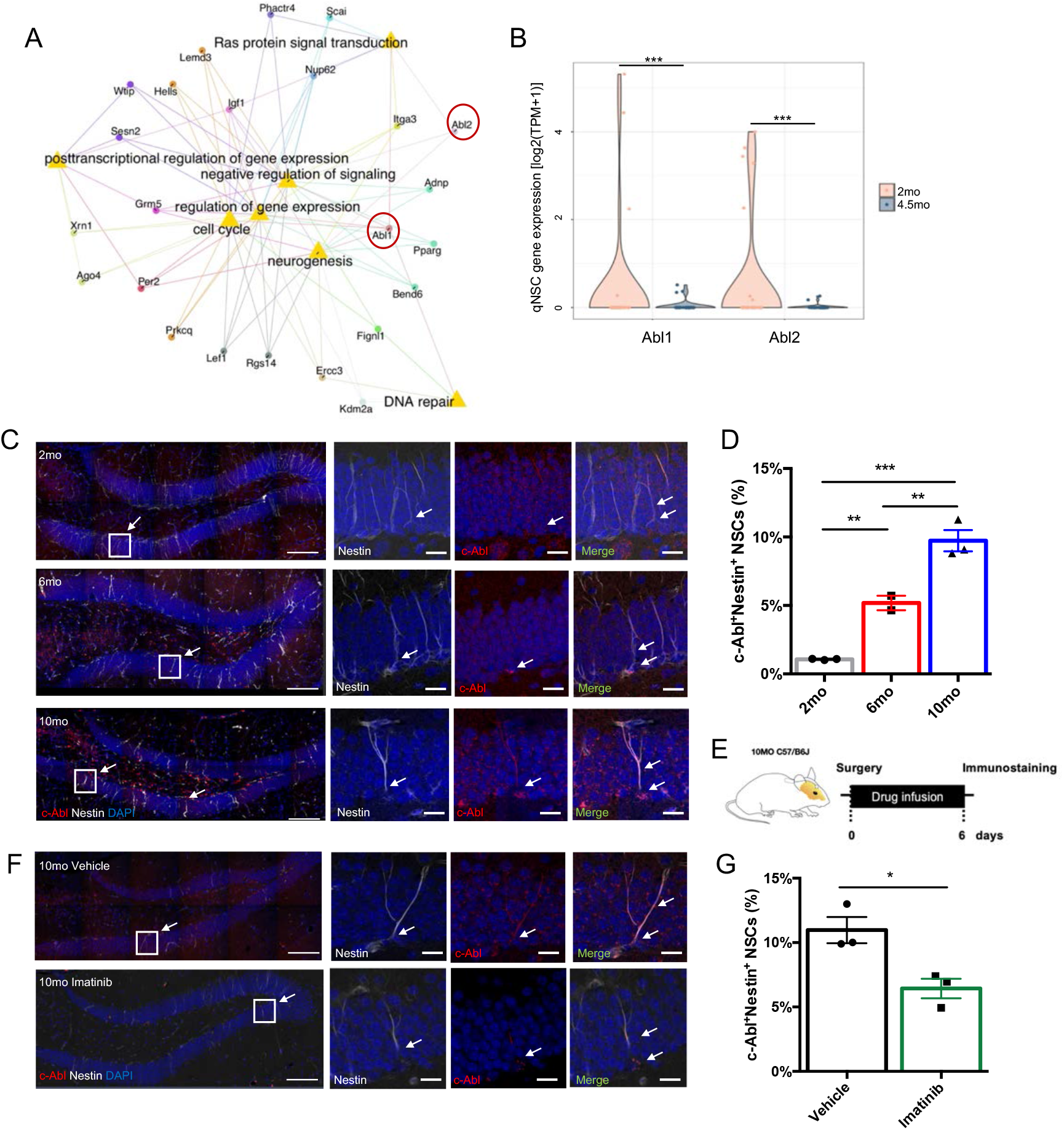
Identification of c-Abl as a NSC aging factor. (A) String network graph depicting age-related changes downregulated in 4.5-month-old qNSCs. Shown genes have 3+ connections to biological processes. (B) Violin plot of Abl1 and Abl2 expression in qNSCs between age groups shows diminished expression of both genes in 4.5-month-old mice. *p<0.05, **p<0.01, ***p<0.001, n.s - not significant; edgeR (quasi-likelihood F-test), FDR-corrected p-values are shown. (C) Representative confocal images of c-Abl and Nestin co-expression across multiple ages (2, 6, 10-month-old). Scale bar, 100μm (20μm for the insert). (D) Quantification of the percentage of double positive NSCs (Nestin^+^ c-Abl^+^) among total Nestin^+^ NSCs in the dentate gyrus of 2-, 6-, 10-month-old mice. Values represent mean ± SEM. (N = 2-3 animals per group; *p<0.05, **p<0.01, ***p<0.001, n.s - not significant; ANOVA with Bonferroni post-hoc test). (E) Schematic illustration of the experimental design for intracranial drug infusion of Imatinib and Vehicle Control. (F) Representative confocal images of c-Abl and Nestin co-expression in Vehicle-Control and Imatinib-treated brains. Scale bar, 100μm (20μm for the insert). (G) Quantification of the percentage of double positive NSCs (Nestin^+^ c-Abl^+^) among total Nestin^+^ NSCs in the dentate gyrus of 10-month-old Vehicle-Control and Imatinib-treated animals. Values represent mean ± SEM. (N = 3 animals per group; *p<0.05, **p<0.01, ***p<0.001, n.s - not significant; ANOVA with Bonferroni post-hoc test).

**Figure 6.**
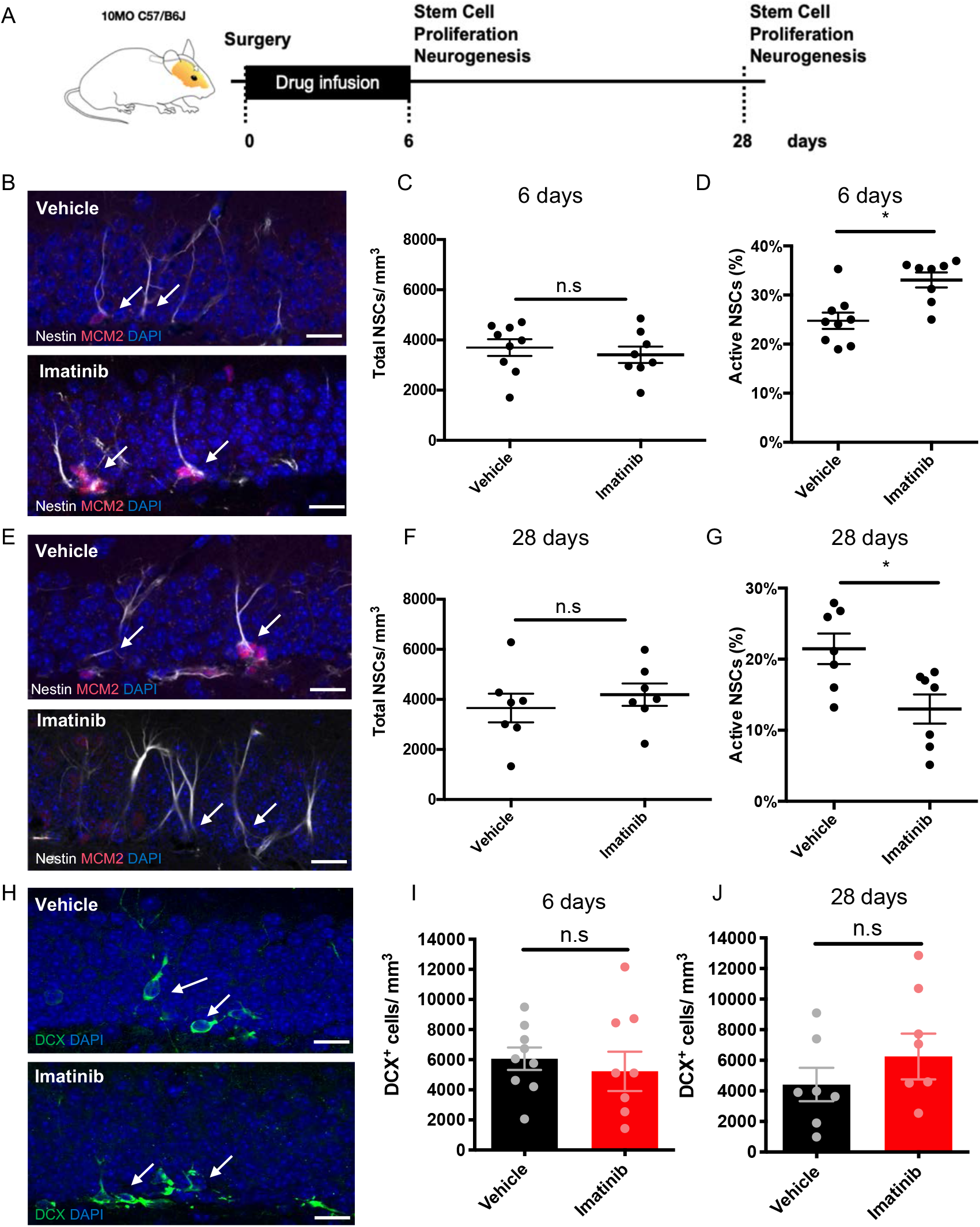
Imatinib partially restores NSC function in the middle-aged brain. (A) Schematic illustration of the experimental design for intracranial drug infusion of Imatinib and Vehicle Control. (B) Sample confocal images showing immunofluorescence for NSCs (Nestin^+^) and cell proliferation (Mcm2^+^) in 10-month-old Vehicle-Control and Imatinib-treated mice for 6 days. Scale bar, 20μm. (C) Quantification of total NSC number (Nestin^+^) in 10-month-old Vehicle-Control and Imatinib-treated mice for 6 days. Values represent mean ± SEM. N = 9 mice (Vehicle), N = 8 mice (Imatinib – treated), *p<0.05, **p<0.01, ***p<0.001, n.s - not significant, Man-Whitney two tailed t-test. (D) Quantification of the percentage of active NSC (Nestin^+^ MCM2^+^) among total Nestin^+^ NSCs in 10-month-old Vehicle-Control and Imatinib-treated mice for 6 days. Values represent mean ± SEM. N = 9 mice (Vehicle), N = 8 mice (Imatinib – treated); *p<0.05, **p<0.01, ***p<0.001, n.s - not significant, Man-Whitney two tailed t-test. (E) Sample confocal images showing immunofluorescence for NSCs (Nestin^+^) and cell proliferation (Mcm2^+^) in 10-month-old Vehicle-Control and Imatinib-treated mice for 6 days and sacrificed at day 28. Scale bar, 20μm. (F) Quantification of total NSC number (Nestin^+^) in 10-month-old Vehicle-Control and Imatinib-treated mice for 6 days and sacrificed at day 28. Values represent mean ± SEM. N = 7 mice (Vehicle), N = 7 (Imatinib-treated); *p<0.05, **p<0.01, ***p<0.001, n.s - not significant, Man-Whitney two tailed t-test. (G) Quantification of the percentage of active NSC (Nestin^+^ MCM2^+^) among total Nestin^+^ NSCs in 10-month-old Vehicle-Control and Imatinib-treated mice for 6 days and sacrificed at day 28. Values represent mean ± SEM. N = 7 mice (Vehicle), N = 7 (Imatinib-treated); *p<0.05, **p<0.01, ***p<0.001, n.s - not significant, Man-Whitney two tailed t-test. (H) Sample confocal images showing immunofluorescence for newborn neurons (DCX^+^) per dentate gyrus section in 10-month-old Vehicle-Control and Imatinib-treated mice. Scale bar, 20μm. (I) Quantification of the number of newborn neurons (DCX^+^) 10-month-old Vehicle-Control and Imatinib-treated mice for 6 days. Values represent mean ± SEM. N = 9 mice (Vehicle), N = 8 mice (Imatinib – treated); *p<0.05, **p<0.01, ***p<0.001, n.s - not significant, Man-Whitney two tailed t-test. (J) Quantification of the number of newborn neurons (DCX^+^) 10-month-old Vehicle-Control and Imatinib-treated mice for 6 days and sacrificed at day 28. Values represent mean ± SEM. N = 7 mice (Vehicle), N = 7 (Imatinib-treated); *p<0.05, **p<0.01, ***p<0.001, n.s - not significant, Man-Whitney two tailed t-test.

The balance between stem cell quiescence and activity is thought to determine the long-term maintenance of the stem cell pool and the neurogenic capacity of the aging brain (Harris et al., 2020; Urbán et al., 2019). Since c-Abl inhibition skews NSC balance toward activation in middle-aged mice, we then sought to determine chronic consequences of short-term Imatinib treatment on NSCs. Imatinib or vehicle control was infused into the hippocampal fimbria of 10-month-old mice for 7 days and mice were sacrificed at day 28. We then assayed total NSC number and proliferation (Nestin^+^MCM2^+^radial glia like cells). Surprisingly, the NSC pool was not prematurely depleted by hyperactivation (Figure 6F), but instead became more quiescent (Figure 6G). Newborn neuron number (DCX^+^ cells) was unchanged in Imatinib-treated brains compared to vehicle control after 6 and 28 days (Figure 6H-J). Together, these data indicate that Imatinib treatment causes NSC activation from a deep quiescence in the middle-aged brain and then NSCs rebound into an even deeper quiescence without prematurely depleting.

## DISCUSSION

Determining when stem cells begin to exhibit age-related dysfunction is a recently recognized approach for identifying targets to slow tissue decline later in life. Here, we show that neural stem cells become compromised early in life within the mature hippocampus. Ascl1^#^-NSCs lack sufficient NSC production and rapidly deplete. Nestin^#^-NSCs were found to temporarily balance self-renewal and differentiation but lose this homeostasis due to an increase in quiescence. NSCs in deep quiescence exhibited changes in cell fate and underwent molecular aging in the mature hippocampus. Importantly, targeting Abl components of this early aging molecular network were able to overcome deep quiescence in middle-aged mice without prematurely depleting remaining NSCs. Finally, we defined a stem cell aging molecular signature shared among neural stem cells, epidermal stem cells, hematopoietic stem cells, and muscle stem cells. These observations suggest that NSCs in the hippocampus are particularly vulnerable to molecular aging, yet their function can be partially recovered by later targeting aging mechanisms.

Neurogenesis is well known to decline with age. Old animals have significantly less NSC numbers, proliferation and neuronal differentiation compared to younger animals (Encinas et al., 2011; Heine et al., 2004; Kuhn et al., 1996; Ziebell et al., 2018). However, heterogeneity within the NSC pool has complicated the association of the cellular origins to age-related neurogenesis decline (Bonaguidi et al., 2011; Dranovsky et al., 2011; Encinas et al., 2011; Pilz et al., 2018). Our single cell lineage tracing suggests that asynchronous NSC dysfunction is a unifying principle of NSC aging. We found that Ascl1^#^-labeled radial cells represent neuronal-biased ST-NSCs, whose number rapidly declines in the young hippocampus. These cells have a higher proliferation rate, primarily generate neurons, and are computationally consistent with “disposable” and “developmental-like” NSC behavior (Encinas et al., 2011; Pilz et al., 2018). Meanwhile, Nestin^#^-labeled radial cells are IT-NSCs that divide and maintain their numbers while ST-NSCs concurrently differentiate. Multipotent IT-NSCs produce neurons, astrocytes, stem cells, and are computationally consistent with “self-renewing” NSCs (Bonaguidi et al., 2011; Dranovsky et al., 2011). However, IT-NSCs increase their time in quiescence with subsequent divisions until homeostasis is compromised and their numbers begin to decline in the mature brain. In addition, IT-NSCs change their fate from asymmetric to symmetric divisions in the mature brain and maintain this lineage bias into middle age. Meanwhile, quiescence continues to increase through middle age. Therefore, hippocampal neurogenesis appears to decline with age in a temporal pattern: (1) ST-NSC number declines in the young brain, (2) IT-NSC quiescence begins to increase in the young brain, (3) IT-NSC number declines in the mature brain, (4) IT-NSC fate changes in the mature brain, and (5) IT-NSC quiescence continues to deepen into the middle-aged brain. This sequence details multiple origins of neurogenesis decline including NSC number, quiescence and cell fate choices (Kuhn et al., 1996, 2018).

While stem cells undergo molecular aging in many old tissues (Goodell and Rando, 2015; López-Otín et al., 2013), only recently have these mechanisms been explored in the brain. Quiescent neural stem cells in the old SVZ exhibit deficits in proteostasis and receive high levels of inflammation (Kalamakis et al., 2019; Leeman et al., 2018). However, transcriptome investigation between young and old SVZ NSCs has thus far not revealed a substantial number of age-related differences. Instead, our clonal lineage in the hippocampus indicated that many changes in NSC function occur in the mature brain. We therefore utilized deeper sequencing to provide enhanced sensitivity (>1000 DEs) of age-related transcriptomic changes in single NSCs. Remarkably, nearly all molecular pillars of cellular aging (Kennedy et al., 2014) were found to change within NSC quiescence. Consistent with NSCs from the aged SVZ (Kalamakis et al., 2019; Leeman et al., 2018; Negredo et al., 2020), NSCs from the mature hippocampus exhibited deficits in proteostasis and received inflammatory signals (NIK-NFkB). Additionally, we uncovered the presence of age-related metabolic alterations, histone demethylase dysregulation, transcriptional downregulation, and DNA damage in NSCs in the mature brain. Our findings strongly implicate that NSCs exhibit early aging in the hippocampus and present associated molecular mechanisms at the systems level.

To determine which molecular pillars of cellular aging are expressed in stem cells more broadly, we sought convergence in transcriptomes from hippocampal NSC, epidermal stem cell, hematopoietic stem cell and muscle stem cells. Similar to prior studies (Keyes and Fuchs, 2018), we did not detect a single age-related gene shared among four stem cells. Instead, we determined common age-related functional deficits in metabolism, RNA homeostasis and proteostasis by gene ontology. Generating string networks to visualize gene-function relationships among 4 stem cells, we observed that gene families such as *Eya, Tet, Fox, Sox, Hox* and *Zfps* are expressed in complementary patterns across individual niches and exhibit biological convergence. These analyses provide a resource that defines intersectional stem cell aging in 2, 3, and 4 stem cell niches. Importantly, we apply this resource to show that the common stem cell chronological aging signature (utilizing MSCs, HSCs, EpSCs from 21-26-month-old mice) is already expressed in NSCs within 4.5-month-old mice. In addition to the intrinsic factors that contribute a large extent to stem cell aging, we also identified changes in Wnt signaling, TGFβ signaling and apoptosis inhibition in 4 stem cell populations. While Wnt and TGFβ family signaling are well known modulators of stem cell activity during aging (Kalamakis et al., 2019; Oh et al., 2014), the role of extrinsic apoptotic signaling inhibition is less studied. Given that extrinsic manipulation of stem cell proliferative homeostasis can extend lifespan (Biteau et al., 2010) as can modulation of metabolism, RNA homeostasis and proteostasis (Chandel et al., 2016; Kim et al., 2018), we provide new targets to potentially coordinate stem cell activity across multiple tissues for treating age-related dysfunction.

Strategies to alter the aging process have recently been developed. Manipulation of blood factors, niche factors, or metabolism (diet and exercise) have demonstrated the capability to partially restore hippocampal neurogenesis and cognition (de Cabo et al., 2014; Horowitz et al., 2020; Katsimpardi and Lledo, 2018; Mahmoudi et al., 2019). In addition, drugs such as rapamycin, metformin, and senoyltics (dasatinib, quercetin) have yielded promising results as potential anti-aging treatments (Leeman et al., 2018; Mahmoudi et al., 2019; Ogrodnik et al., 2019; Torres-Pérez et al., 2015). However, the vast majority of these studies have been performed in animals of advanced age (Torres-Pérez et al., 2015). Instead, age-related changes to physiology accumulate early in life, and interventions to reverse or delay age-related diseases are more effective earlier in disease etiology (Belsky et al., 2015; Gillman, 2005). We have taken a similar approach by investigating the initiation of NSC aging and functional decline in the mature brain. Importantly, targeting molecular aging with Imatinib in the middle-aged brain was sufficient to overcome deep NSC quiescence. In addition, NSC number is maintained through this process, which is unexpected given the observations of a finite NSC pool containing limited self-renewal capability (Ehm et al., 2010; Encinas et al., 2011; Jones et al., 2015; Mu and Gage, 2011; Pilz et al., 2018; Renault et al., 2009; Sierra et al., 2015). We provide, to the best of our knowledge, the first evidence of Abl signaling directly regulating NSC biology and its role as an NSC aging factor. Since increased quiescence and resistance to enter cell-cycle occur in many old tissues that exhibit stem cell dysfunction (Fan et al., 2017; Goodell and Rando, 2015; Kalamakis et al., 2019), this approach may provide targets for early interventions in multiple tissues. However, imatinib does not fully “rejuvenate” NSCs because after induced cell cycle, NSCs enter a deeper quiescence and do not increase neurogenesis typical of younger NSCs. Therefore, understanding the key mechanisms that drive stem cell aging will help to create new directions towards age-related regenerative medicine.

## ACKNOWLEDGMENTS

We thank David Cobrinik, Sunhye Lee and members of the Bonaguidi, Simons and Jang laboratories for invaluable discussions and comments on the manuscript; Teresa Krieger, Dennisse Jimenez-Cyrus, Avinash Iyer and Lee McMahon for technical support; CHLA FACS Core Facility, CHLA Center for Personalized Medicine Sequencing facility for technical assistance. This work was supported by grants to M.A.B. from the NIH (R00NS080913, R56AG064077), Donald E. and Delia B. Baxter Foundation, L.K. Whittier Foundation, Eli and Edythe Broad Foundation grants and CHLA TSRI pilot program; M-H.J. was supported by the NIH (RO1AG0585560); B.D.S. was supported by a Royal Society E P Abraham Professorship (RP\R1\180165) and Wellcome Trust (grant number 098357/Z/12/Z); A.I. was partially supported by the American Federation for Aging Research Scholarship for Research in Biology of Aging and NIH T32HD060549-08 fellowship, and M.B. was partially supported by the USC Provost’s Research Enhancement Fellowship.

## AUTHOR CONTRIBUTIONS

Conceptualization: A.I., M.B. and M.A.B.; Data collection and methodology: A.I., E.P., L.P., N.Z., H.J., M.A.B., and C.L, G.R, A.H. with supervision; Data analysis and bioinformatics: A.I., M.B., D.J, L.P., D.A., B.S. and M.A.B.; Resources: M-H.J, B.S, M.A.B, Supervision: M-H.J, B.S, M.A.B.; Writing, Review and Editing: A.I., M.B. and M.A.B.; Project coordination: M.A.B. All authors commented on the manuscript.

## STAR★Methods

### Key Resource Table

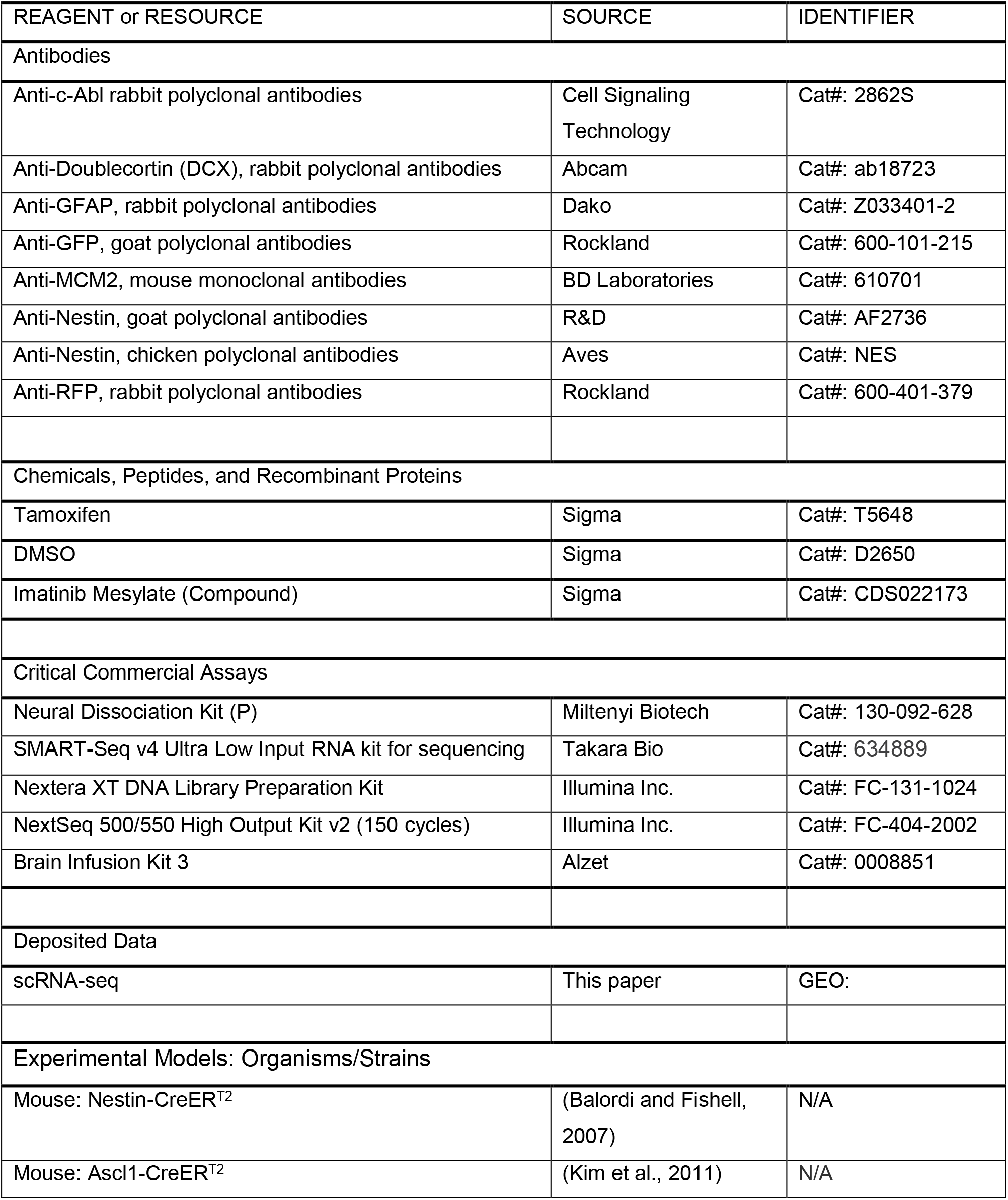
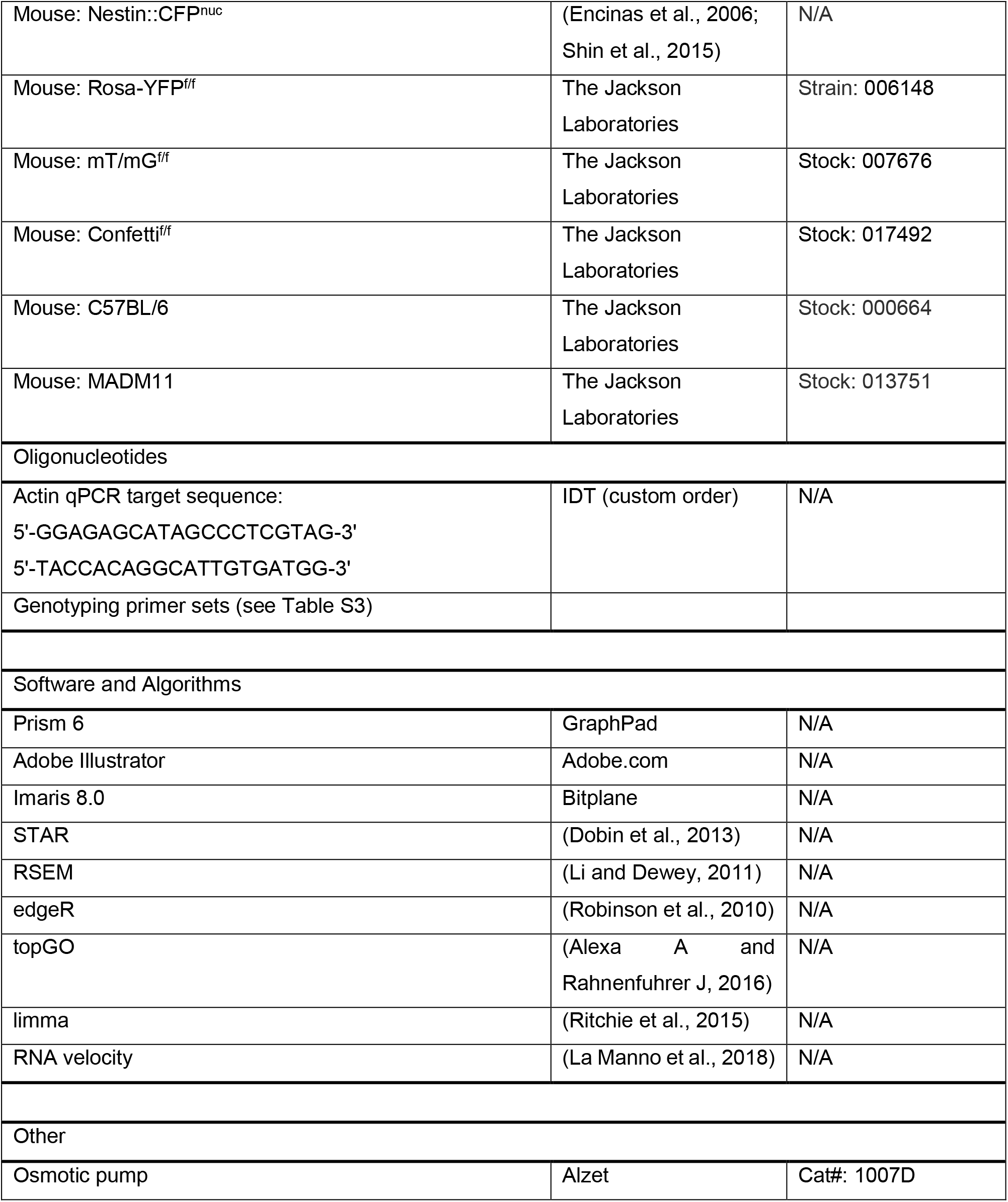

#### Contact for Reagent and Resource Sharing

Further information and requests for resources and reagents should be directed to and will be fulfilled by the Lead Contact: Michael A. Bonaguidi

### Experimental Model and Subject Details

#### Animals and Tamoxifen Administration

All animal procedures were performed in accordance with institutional guidelines of University of Southern California Keck School of Medicine and protocol (20287) approved by Institutional Animal Care and Use Committee (IACUC). All mice used in the study were backcrossed to the C57BL/6 background to ensure the reproducibility of clonal induction with specific doses of tamoxifen. Animals were housed in a 12-hour light/12-hour dark cycle with free access to food.

Nestin-CreER^T2^ mice (Balordi and Fishell, 2007) and Ascl1-CreER^T2^ mice (Kim et al., 2011) were used to clonally label RGLs. The following genetically modified mice were originally purchased from Jackson Labs: Rosa-YFP^f/f^ (Strain: B6.129X1-Gt(ROSA)^26Sortm1(EYFP)Cos^/J), mT/mG^f/f^ (Strain: B6.129(Cg)-Gt(ROSA)^26Sortm4(ACTB-tdTomato^,^−EGFP)Luo^/j), Confetti^f/f^ (StrainGt(ROSA)26Sor^tm1(CAG-Brainbow2.1)Cle^/J).

Nestin-CreER^T2^ and Ascl1-CreER^T2^ mice were crossed to fluorescent reporter mice for clonal analysis. Nestin-CreER^T2^::Confetti^f/+^ mice were generated by breeding Nestin-CreERT2+/- mice with Confetti^f/f^ mice, or by crossing Nestin-CreER^T2+/-^::Confetti^f/f^ mice with wild-type C57BL/6 mice. Ascl-CreER^T2+/-^::Confetti^f/f^, Ascl1-CreER^T2+/-^::YFP^f/+^or Ascl1-CreER^T2+/-^::mTmG^f/+^ mice were generated by crossing Ascl1-CreER^T2+/-^ with Confetti^f/f^, YFP^f/f^ or mTmG^f/f^, respectively. For each CreER^T2^ driver, we tested different tamoxifen doses and reporter lines, and obtained combinations that exhibited high specificity, inducibility and reproducibility (Table S1, S3–S4).

At least 3 animals were checked for each reporter/driver combination to ensure there was no recombination in the adult SGZ in the absence of tamoxifen. A stock of tamoxifen (66.6 mg/mL) was prepared in a 5:1 ratio of corn oil to ethanol at 37 °C with occasional vortexing. A single tamoxifen or vehicle dose was intraperitoneally injected into 8-to 10-week-old, 26-to 30-week-old, or 56-week-old mice at various concentrations for lineage tracing (Table S1). Injected animals showed no signs of distress.

Primer sets (Table S3) from original publications were used to identify genetically modified mice (Ahn and Joyner, 2005; Balordi and Fishell, 2007; Kim et al., 2011; Lemberger et al., 2007; Muzumdar et al., 2007). Genomic tail DNA was isolated in a 25mM NaOH, 0.2 mM EDTA solution and ran for 35 PCR cycles.

### Method details

#### Immunostaining, Confocal Imaging, and Processing

Mice were anesthetized with isoflurane gas and underwent transcardial perfusion with saline followed by 4% paraformaldehyde. Brains were post-fixed overnight in 4% paraformaldehyde and then immersed in 30% sucrose for a subsequent 48 hours prior to sectioning. Brains were sectioned into 45μm coronal sections through the entire dentate gyrus. Immunohistology was performed with antibodies as previously described (Bonaguidi et al., 2011) on sections in serial order using custom, in-house staining chambers. Brain sections were washed in TBS with 0.3% Triton-X100 prior to staining and mounting. Goat anti-GFP (1:1000), rabbit anti-RFP (1:1000), rabbit anti-GFAP (1:2000), chicken anti-Nestin (1:500), mouse anti-MCM2 (1:500), rabbit c-Abl (1:75), rabbit Dcx (1:500) primary antibodies were used. Antigen retrieval for MCM2, Nestin and c-Abl antibodies utilized DAKO citrate buffer (Dako, Cat#: S1699) at 95°C for 30 min and then samples were left for 1 hour to cool at room temperature. Cells were then counted using every 8th section throughout the entire dentate (n = 6-7 sections per dentate) for a stereological analysis and every section was processed for clonal analysis. Fluorescent-labeled cells were identified with a Zeiss AxioObserver.A1 microscope and were acquired as a z-stack on a Zeiss LSM700 confocal system under 40X or 63X magnification. Morphological analysis was done using Imaris 8.0 Software. Radial-glial like neural stem cells were determined by the presence of Nestin or GFAP protein or in a radial direction into the granule cell layer. Co-expression with Mcm2 was assessed by a Nestin ring or GFAP “y-shape” around DAPI-labeled nuclei for the presence of Mcm2.

#### Clonal Analysis

Clonal analysis and categorization of cell types by morphological and immunohistological criteria were consistent with prior criteria (Bonaguidi et al., 2011). Analyzed dentate volume included the stratum granulosum (granule cell layer), and SGZ. Serial sections were first screened for candidate clones, which were defined as possessing at least (1) an NSC (RGL), (2) neuronal cell(s) in close spatial proximity, or (3) astroglia in close spatial proximity to other astroglia or neuronal cells. Approximately 6 - 24 clones per dentate per color (in the case of Confetti) allowed for clonal analysis based on prior computer simulations (Bonaguidi et al., 2011) (Table S4). Clones were randomly induced throughout the dentate.

Dividing clones were marked with two separate nuclei by DAPI under confocal microscopy. The two nuclei were completely encompassed when using membrane-bound reporters (Rosa-CFP, mT/mG, Confetti-CFP), or completely filled when using cytoplasmic-bound reporters (Rosa-YFP, Rosa-RFP, Confetti-YFP, Confetti-RFP). Clones were categorized according to the clone composition among RGL-containing clones (Bonaguidi et al., 2011). Clones with more than one NSC and progeny were classified by the lineage produced: neuronal, astroglial, or both. NSC maintenance was assessed as a percentage of clones that contained at least one NSC.

#### Stereotaxic surgery (Imatinib infusion via osmotic pump)

Stereotaxic surgery was performed on 10-months-old C57/BL6 mice as described (Pan, 2015; Pan et al., 2012; Wong et al., 2000). Mice were anesthetized using an isoflurane machine (5% until recumbent, 2-3% maintenance). Prior to anesthesia, nonsteroidal anti-inflammatory analgesic - 1X Ketoprofen (5 mg/kg) was injected. Osmotic pump with the drug (1 mM Imatinib treated) or 1 % DMSO (Vehicle control) was unilaterally implanted at a rate of 0.5μl/hr for 6 days (Figure 5 and 6) into the hippocampal fimbria with the following coordinates relative to Bregma: - 0.8 mm posterior, - 0.75 mm medial-lateral, and 2.5 mm ventral and mice were sacrificed on the day 7 or day 28 after drug infusion.

#### Fluorescence-activated Cell Sorting (FACs) of Individual cells from Adult Mouse Dentate Gyrus

Homozygous Nestin:CFP^nuc^ mice (Encinas et al., 2006) were used for all singlecell RNA-seq experiments (Shin et al., 2015). Mice were euthanized by cervical dislocation, and brains were immediately immersed into cold Hibernate A solution (BrainBits). Dentate gyri were dissected under a stereomicroscope as previously described (Hagihara et al., 2009). All procedures were performed with approved protocols in accordance with institutional animal guidelines. The single cell suspension was prepared by using Neural Tissue Dissociation Kit (P), with the addition of one cleaning step with Percoll (1:10 dilution) to remove myelin layer and cellular debris. Propidium Iodide was added to determine cell viability and cells were sorted on a BD FACs Aria II with a 70 μm nozzle at 13 psi (Figure S3). Single cells were collected onto the ice-cold 8-sample parafilm-covered glass slides with 1.25uL of 1X lysis mixture (Clontech Laboratories SMART-Seq v4 Ultra Low Input RNA Kit for sequencing) and immediately transferred in individual 0.2-ml RNase-free 8-well strip (Figure S3). To avoid any effects due to the FACs sorting protocol, cells were kept on ice the entire procedure, and the actual collection was performed at 4°C.

#### scRNA-seq library preparation and sequencing

cDNA was generated using the SMART-Seq v4 Ultra Low Input RNA Kit (Clontech Laboratories) for sequencing with modifications to bring down the cost. Briefly, all reagents from the kit (for Reverse transcription and cDNA amplification) were miniaturized 10-fold by utilizing FACs collection in a very low volume of the lysis buffer. The same quality of cDNA was achieved as for the original kit. The amplification product was purified using Ampure XP beads according to the manufacturer’s protocol. Quality control and validation were performed by using qPCR for Actin expression and Agilent TapeStation 4200 (Agilent Technologies).

The amplification product was sheared using library preparation Nextera XT Kit (Illumina Inc.) following the manufacturer’s protocol. In brief, the purified cDNA was tagmented and fragmented using a miniaturized protocol. cDNA fragments were barcoded using i5 and i7 indices (Illumina Inc.) and the product was purified using Ampure RNA clean beads. The purified ligation product underwent 13 cycles of PCR. A Mantis Liquid Handler (Formulatrix, Inc.) was used to handle low volume pipetting. The libraries were multiplexed and sequenced using Illumina NextSeq 550 (Illumina Inc.) 75 paired end run. To minimize the batch effect on the sequencing all cells were loaded into one chip.

#### Preprocessing, Alignment and Count Estimation of RNA-Seq data

FASTQ files were demultiplexed into two cell-specific FASTQ files by i5 and i7 adapter sequences for forward and reverse ends of the paired-end reads. Raw reads were trimmed of TSO sequence and adaptor contaminants using Trimmomatic (Bolger et al., 2014). Sequences were then aligned using STAR (Dobin et al., 2013) to the mm10 mouse transcriptome, constructed using UCSC annotation framework, generating cellspecific BAM files. Only uniquely mapped reads, concordant between both directions of the paired end reads were preserved. RSEM (Li and Dewey, 2011) was then employed to produce spliceoform-level count estimates, which were summated to give gene-level counts per cell. Cells with fewer than 150,000 uniquely mapped reads were excluded. Remaining cells then underwent transcripts per million (TPM) normalization (Li and Dewey, 2011) to control for variable gene length within samples and sequencing depth between samples. Cells with fewer than 1,000 detected genes were also excluded, where a gene was considered detected in a cell if > 2 TPM. After cell exclusions, 48 cells from 2-month-old mice and 41 cells from 4.5-month-old mice remained, totaling 89 cells. 3,603 genes were detected on average (SD = 833).

#### Clustering, Dimensionality Reduction and RNA Velocity

K-means clustering, employing the Hartigan-Wong algorithm, was run on all 89 cells in full 24,411-dimensional expression space. K was set equal to 5, as (Shin et al., 2015) described 5 clusters present in adult neural stem cells. Dimensionality reduction of 89 cells was performed by PCA, using the FactoMineR (Lê et al., 2008) package with default parameters.

RNA Velocity (La Manno et al., 2018) was used to estimate the future expression state of single cells. Count estimates of unspliced and spliced cells were computed with the following command *velocityto run_smartseq2-d 1 $bam_files $ucsd_annotation_mm10.gtf*. Unspliced and spliced count estimates were imported to the gene.relative.velocity.estimate() function to produce future expression state estimates for every cell. Future expression states were represented in previously computed PCA embedding as arrows. An independent samples t-test was conducted to compare the magnitude of future state prediction vectors from 2 month old and 4.5 month old individual cells (Figure 3F). The cells from 4.5 month old animals had significantly longer vectors in principal component space (M = 4,568.99, SD = 2,095.80) than cells from 2 month old animals (M = 3,292.23, SD = 1,589.42); t(49.94) = 2.63, p = 0.011.

To represent the extent to which aging dampens the tendency for cells to proceed down the neurogenic trajectory, we visualized the future state vectors of each age group at each cluster. Vectors were centered on the mean coordinate within their respective cluster in PC space. The X corresponds to PC1 coordinate space, where Y corresponds to PC2 coordinate space. First mean, coordinates for each cluster were derived. Cells within any one cluster were computed thusly –

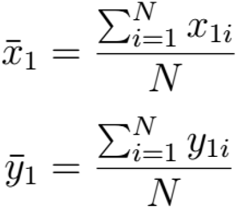

Future state predictions were then aligned atop each other at the mean coordinate by -

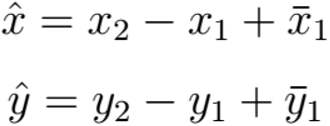

This computation was run independently on cells from the 2-month-old and 4.5-month-old groups to produce single cell estimates.

#### Differential expression, niche comparisons and Gene Ontology enrichment

Differential expression on NestinCFP^+^ scRNA-Seq generated for this paper was computed using the edgeR package in R (Robinson et al., 2010). Specifically, we employed the general liner model quasi likelihood fit test edgeR (edgeRQLFDetRate), as it was optimal across all major evaluation criteria in a recent survey of contemporary differential expression methods developed for single cell RNA Sequencing (Soneson and Robinson, 2018). In all use cases, age group was the only factor for the generalized linear model, where there were two age groups (2-month-old and 4.5-month-old). Gene filtering is required for accurate implementation of edgeR. Only genes with a mean count greater than 5 in at least one of the age groups were included any analysis. P-values were group-adjusted using the Benjamini-Hochberg (Hochberg and Benjamini, 1990) method to produce a false detection rate (FDR). Genes with an FDR <=0.05 were preserved for downstream analyses.

Transcriptomes from epidermal stem cell (Solanas et al., 2017) and muscle stem cell (Liu et al., 2013) isolations were observed on Affymetrix Mouse Gene microarrays. Both datasets were obtained via GEOquery (Sean and Meltzer, 2007) using the getGEO() function. To obtain differentially expressed gene sets between young and aged cells, samples were first log transformed and quantile normalized. Expression units were then fit to a linear model where age was the single factor. Finally, an empirical Bayes statistical test was run on each bead using default parameters of the eBayes() function in the limma package (Ritchie et al., 2015). P-values were group-adjusted using the Benjamini-Hochberg (Hochberg and Benjamini, 1990) method to produce a false detection rate (FDR). As some beads encompassed multiple genes, gene ID names were disaggregated to create a differentially expressed gene set of all gene names implicated by the empirical Bayes test. Differentially expressed genes between young and aged hematopoietic stem cells were obtained from tests run by (Sun et al., 2014).

To determine if the convergence between the early aging expression signature observed in NSCs and the more advanced aging signature of other somatic stem cells was statistically significant, we ran a Fisher’s exact test (Fisher, 1956) comparing the genes differentially upregulated in NSC aging against the genes differentially upregulated with aging in each of the other stem cell niches against a gene universe encompassing genes jointly present on the Affymetrix microarray and UCSC genome annotation. The same series of Fisher’s tests was run on downregulated genes.

TopGO (Alexa et al., 2006) was employed to derive enriched Gene Ontology terms associated with aging in each stem cell niche. Two ontology enrichment lists were generated for each niche - one from upregulated genes and another from downregulated genes. Enrichment was calculated using the classic Fisher’s test (Fisher, 1956), where exact p-values were corrected for multiple comparisons using Benjimini-Hochberg (Hochberg and Benjamini, 1990).

As was performed at the level of genes, Gene Ontology (GO)-level convergence in aging signatures between NSCs and other stem cell niches was also examined. A Fisher’s exact test was employed between upregulated NSC GO terms and the upregulated GO terms in each of the other niches. Downregulated terms were similarly examined.

SuperExactTest (Wang, Zhao and Zhang, 2015) was performed on the intersection of each combination of DE gene sets and enriched GO term sets for all four stem cell niches. Importantly, the probability that all four stem cell niches converged onto 18 or more common GO terms is highly unlikely under random selection (p = 6.99×10^−77^) (Figure 4D-E).

#### String network visualization

To represent connectivity between genes and GO terms associated with NSC aging, we wrote custom functions to visualize a string network. The underlying object for this network is represented as the bipartite graph *G* = (V,E) where, V(*G*) is a set of vertices, which can be partitioned into two independent sets, GO terms A, and genes B. To reduce visual complexity, we generated the subgraph *H* where V(*H*)⊆ V(*G*) and *E*(*H*)⊆ E(). The minimum allowable edges is *a*, thus conditionally *δ(H) = a*. Construction of graphs connecting genes to both GO terms and niches is tripartite yet follows the same design as the previous bipartite graph, with the addition of a new subset to V(*G*). Once constructed, graphs were matricized and visualized as a string network using force-directed Kamadakawai spatial organization in the GGally package (Schloerke et al., 2014).

### Computational modeling

To consolidate the discussion in the main text, in the following we further explain the fitting and modelling approaches used.

#### Homeostasis

Homeostasis is defined as the situation in which the average stem cell number per clone remains constant over time, taking the value unity if only stem cells are labelled at the time of induction. Homeostasis was determined at each time point by quantifying the average number of radial glial-like NSC per all clones (Figure 1L). NSC clonal maintenance (Figure 2B) was assessed by quantifying the proportion of clones which contains NSCs among all clones. NSC self-renewal fits were used to predict the number of NSCs per NSC-retaining clone required to offset NSC clonal depletion (Figure 2B) as a function of time (age). We performed a nonlinear regression - second order polynomial - to fit the Nestin# data and a scatter nearest neighbor to fit Ascl1# data to determine the predicted value (PV) curves in Figure 2C.

#### Distribution of activation times

For Figure 2F, the activation time of a radial glial-like NSC (RGL) is the time taken to first enter into cell cycle upon labelling (*T_entry_*). To deduce the distribution of activation times from the clonal data, we calculated at each time point the fraction of RGLs that had not yet divided.

As the labelling protocols targeted a small proportion of IPCs in addition to RGLs, there was a degree of error involved in our assignment: we could not decide unambiguously whether a clone consisting entirely of IPCs and differentiated cell types was originally derived from an RGL or an IPC. In the case of Ascl1#-NSC clonal data, the number of such clones was so small that their exclusion would not significantly affect our results. For Nestin#-NSC, 10/47 clones consist only of 1-2 IPCs at 2 dpi; we assumed that these were IPC-derived and excluded them from the analysis. By 7 dpi, due to the short cell cycle time of IPCs (Berg et al., 2018; Hayes and Nowakowski, 2002; Hodge et al., 2008), we would expect IPC-derived clones to have grown in size; we therefore took clones containing 6 or more IPCs to be IPC-derived (2/40), and clones consisting only of 2 IPCs to be RGL-derived (4/40). At later time points, we assumed that all clones were RGL-derived. As a second caveat, a clone consisting of a single RGL was scored as an undivided RGL but may have given rise to progeny that were subsequently lost through cell death. However, effects of any erroneous assignments were likely negligible compared to experimental noise over this timescale.

By performing a weighted least-squares fit, we deduced that Nestin#-NSCs enters cell cycle at a rate = 0.044 ± 0.005 per day, equivalent to a mean cell cycle time of = 23 ± 3 days. Similar fits to the early-time Ascl1-CreER^T2^ data suggested mean activation times of 0.35 ± 0.04 days, respectively.

#### Cell cycle re-entry times

For Figure 2G we can estimate cell cycle re-entry times from the fraction of cells that have divided exactly once. For a population of cells with activation rate *λ* and cell cycle re-entry rate *μ*, the fraction of cells divided once at time *t*, denoted *R*^1^(*t*), satisfies

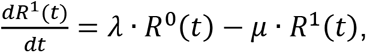

where *R*^0^(*t*) = *e^λt^* is the fraction of cells that remain undivided at time *t*.

Multiplying by the integrating factor *e^μt^* yields

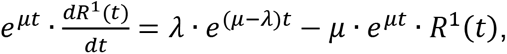

with the solution 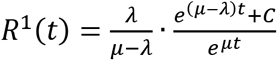, where *C* is a constant. Since *R*^1^(0) = 0, we must have *C* = −1 and therefore 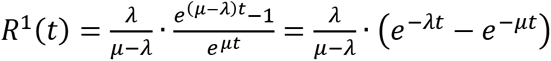.

Given our estimates of the activation times, weighted least-squared fits suggested cell time to the first Nestin# - NSC division for 2 months old is 23 ± 3 days, for 6 months old is 38 ± 3 days; cycle re-entry times for 2 months old is 40 ± 3 and for 6 months old is 100 ± 4 days (Figure 2E-G).

#### Model of a developmental-like stem cell fate program (Short Term - Ascl1#-NSC)

Here we introduce a simplified Markovian model that captures the progression of radialglia like neural stem cells (NSCs) through a developmental-like programme that consists of a proliferative phase, followed by a neurogenic phase and terminal differentiation (Pilz et al., 2018). In our model, NSCs divide with a transition rate *λ* (corresponding to an average cycle time *T* = *λ*^−1^) and go through *R*_0_ rounds of duplication (symmetric self-renewing divisions), followed by *R*_1_ rounds of asymmetric division, generating a certain amount of neuronal progeny upon each division, followed by *R*_2_ = 1 round of terminal differentiation. During each differentiating division, the amount of neuronal progeny generated (interpreted as containing all cell types down the neuronal lineage) is exponentially distributed with a mean proliferative output *m_N_*. This procedure takes into account amplification by intermediate progenitors. Neuronal progeny dies at a rate *ω_N_*. Initially, a clone starts as a single NSC (with probability *q*) with uniform probability to find it at any state within the programme or a neuronal progenitor/neuron (with probability 1 − q).

Formally, the state of a clone is given by the vector (*r,n*) = (*r*_0,0_,…,*r*_0,*R*_0_−1_,*r*_0,1_,…,*r*_0,*R*_1_−1_,*r*_2,0_,*n*), where *r_σi_* is the number of NSCs that are in phase *σ*, where *σ* = 0 corresponds to the proliferative phase, *σ* = 1 corresponds to the phase of asymmetric neurogenic divisions and *σ* = 2 corresponds to the phase of terminal differentiation, and *i*. is the number of divisions that a cell has gone through in the respective phase; *n* denotes the neuronal content. The dynamics of the probability *P* = *P*(*r,n,t*) to find a clone with configuration (*r,n*) at time t is described by the master equation

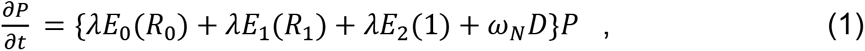

where we have defined the following operators: *E_σ_*(*R*) describes the progression through the phase *σ* = 0,1,2 of the developmental-like programme,

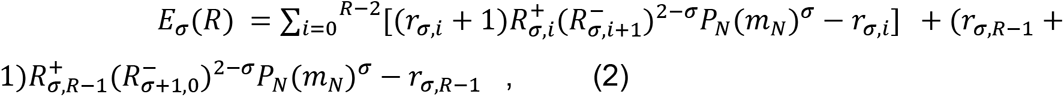

where the 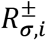 are ladder operators that increase/decrease the number of RGLs at stage (*σ,i*) according to 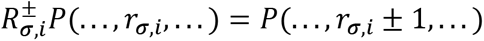 and

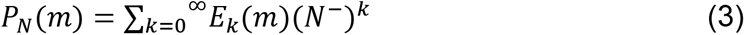

is the operator that generates neuronal progeny with an exponentially distributed number of newborn cells with average *m* with *E* being a discrete exponential distribution; here, *N*^±^ are the ladder operators for the variable *n*. Furthermore, we define the operator describing death of neuronal progeny as

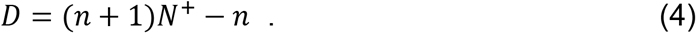

Here, we impose the boundary condition *P*(*r, n,t*) = 0 if *r_σi_* < 0 for at least one (*σ, i*). The initial condition described above is formally given by *P*(*r, n,* 0) = *q*(*R*_0_ + *R*_1_ + *R*_2_)^−1^ + (1 − q)*δ_n,1_δ_r_σ,i__*.

Motivated by the fact that Ascl1#-targeted cells seem to be primed for entry into cycle (see main text), we allow the first division to occur at a faster rate *λ*_1_ = *αλ*, where *α* > 1 indicates the fold-change as compared to the re-entry rate *λ*. Formally, this entails a straightforward modification of Eq. (1), which we omit here since it obscures the basic features of the stochastic dynamics. The model is solved numerically using a standard stochastic simulation algorithm of the Gillespie-type (Gillespie, 1977).

To describe the clonal dynamics of the Ascl1^#^-targeted population, we identify model parameters using the following strategy. First, all parameters are determined simultaneously by comparing the model to the 2-months age dataset, fitting the timedependent average clone content 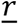, the time-dependent ‘RGL survival’ 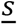, defined as fraction of clones containing at least one RGL (Figure S2) and the average number 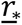 of RGLs per RGL-containing clone,

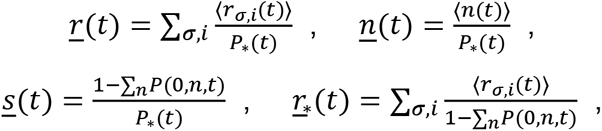

where *P*_*_(*t*) = 1 − *P*(0,0,*t*) is the total surviving fraction of clones. For successive later ages, all parameters are held constant except for the neuronal death rate *ω_N_*, which is determined individually for each age by refitting the model to the respective dataset. To implement this fit strategy, we compute the sum of squared differences between experimental data points and simulations for each of these quantitites individually and then obtain a combined cost function *R* by multiplying these individual residuals,

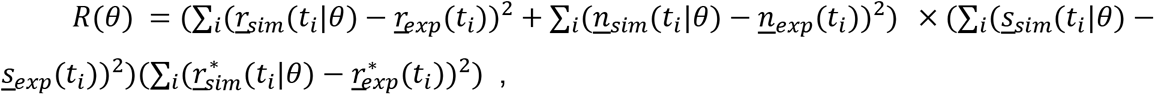

where *θ* is the respective set of parameters and *t_i_* are the experimentally available time points. The cost function *R* is minimized using a stochastic optimization algorithm (a Covariance Matrix Adaptation Evolution Strategy using the cma-es Python package (Igel et al., 2006). All fit parameters are found in Figure S2.

#### Model of cell fate dynamics within a small stem cell niche

In this model, NSCs divide within a small niche region that can harbor a maximum of *N* NSCs. NSCs stochastically divide at a rate *λ*. If the niche is not maximally occupied, NSCs duplicate with probability *p* and differentiate with probability 1 − *p* upon which NSCs are lost while generating a certain amount of neuronal and glial progeny. As in the model of the developmental-like programme, the amount of progeny generated (interpreted as containing all cell types down the neuronal or glial lineage, respectively) is exponentially distributed with a mean proliferative output *m_N_* or *m_G_*, respectively. If the niche is maximally occupied, the duplication channel is blocked, and cells differentiate with probability 1. Neuronal and glial progeny dies at rates *ω_N_* and *ω_G_*, respectively. The state of a clone is given by the vector (*r,n,g*) where *r* is the number of NSCs, *n* denotes the neuronal content and *g* denotes the glial content. Initially, the clone starts as a single NSC. The dynamics of the probability *P* = *P*(*r,n,g,t*) to find a clone with configuration (*r,n,g*) at time *t* is described by the master equation

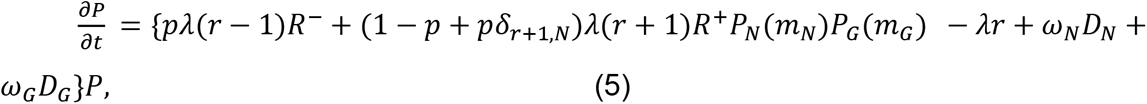

where the operators *P_x_* and *D_x_* being defined analogously to Eqs. (3) and (4) with *P_G_* given by *P_N_* with *N*^±^ being replaced by the ladder operator *G*^±^ for the glial progeny number *g*. Here we impose the boundary conditions *P*(−1,*t*) = *P*(*N* + 1,*t*) = 0, which also ensure that the niche occupation cannot exceed its maximum capacity *N*. As for the developmental-like model, we add the possibility for the first division to occur at a faster rate *λ*_1_ = *αλ*.

Parameters are determined using the following strategy. First, all parameters are determined in the same way described in the previous section for the developmental-like programme (Figure S2). Cycle times at later ages are inferred from the total population data, an independent data set (Figure S2). To this end, we fit the total RGL volume *V*(*t*) by a function of the form *V*(*t*) = *V*_0_ + *V*_1_*e^−kt^* and assume that the time evolution of the RGL loss rate *η*, defined by *V*, = −*ηV*, reflects changes in cycle time. Hence, we infer relative changes in the cycle time at later age by the factor

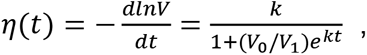

so that

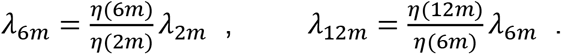

Using the thus inferred cycle rates and the other fit parameters, only the neuronal and glial death rates are determined using refits at 6 months of age. All fit parameters are found in Table S2.

### Statistics

Statistics were performed as indicated in each Figure legends. For *in vivo* experiments, the number of independent experimental replicates are indicated in figure legends, with n representing n experimental replicates (clones) using at least n animals. For a single cell RNA-seq experiment, 2-3 mice per group were used to obtain enough cells. Mann-Whitney unpaired two tailed t-test was conducted to compare the Imatinib effect against Vehicle controls. All multiple comparisons were performed with GraphPad Prism’s one-way ANOVA function with Bonferroni’s multiple comparisons test. All clones observed at each time point were treated as statistically equivalent. No randomization or blinding was used in the animal studies. Mouse dentate gyri with exceedingly low (<6) or high (>24) clones per color (Confetti) per hemisphere were omitted from the study. Sample sizes were estimated in accordance to prior clonal studies (Bonaguidi et al., 2011; Song et al., 2012). Error bars in the study represent the standard errors in mean frequencies, calculated as ✓(p(1-p)/N) where p is the frequency of a given characteristic and N the number of clones considered. Mathematical modeling was performed by weighted least squares using custom-written MATLAB scripts. Representative images are depicted in Figures: 1, S1–S3.

## Supplementary Tables

**Table S1.** Tamoxifen doses used to achieve clonal recombination among various promoter, reporter and ages contexts over the analyzed time course. (**Total number of clones acquired = 1561, total number of animals used = 130).**

| Promoter | Reporter | Age (month-old) | [Tamoxifen] | Chase | Number of clones | Figure(s) |
| --- | --- | --- | --- | --- | --- | --- |
| Ascl1::CreER<sup>T2</sup> | ROSA-YFP | 2 | 78 mg/kg | 1, 3, 7, 30, 60, 120 dpi | 1dpi = 40<br>3dpi = 83<br>7dpi = 92<br>30dpi = 70<br>60dpi = 59<br>120dpi = 80 | 1, S1 |
| Ascl1::CreER<sup>T2</sup> | mT/mG | 2 | 78 mg/kg | 1 dpi | 1dpi = 45 | 1, S1 |
| Ascl1::CreER<sup>T2</sup> | mT/mG | 6 | 233.1 mg/kg | 3, 7, 30 dpi | 3 dpi = 63<br>7dpi = 101<br>30dpi = 78 | 1,2, S1-2 |
| Ascl1::CreER<sup>T2</sup> | Confetti | 6 | 233.1 mg/kg | 1 dpi | 1 dpi = 30 | 1,2, S1-2 |
| Ascl1::CreER<sup>T2</sup> | mT/mG | 12 | 233.1 mg/kg | 7, 30 dpi | 7dpi = 52<br>30 dpi = 69 | 1,2, S1-2 |
| Ascl1::CreER<sup>T2</sup> | ROSA-YFP | 12 | 233.1 mg/kg | 30 dpi | 30 dpi = 35 | 1,2, S1-2 |
| Nestin::CreER<sup>T2</sup> | Z/EG | 2 | 62 mg/kg | 2, 7, 30, 60, 120 dpi | 2dpi = 34<br>7dpi = 40<br>30dpi = 42<br>60dpi = 45<br>120dpi = 69<br>365dpi = 30 | 1, S1 |
| Nestin::CreER<sup>T2</sup> | MADM | 2 | 62 mg/kg | 2, 60 dpi | 2dpi = 13<br>60dpi = 24 | 1, S1 |
| Nestin::CreER<sup>T2</sup> | Confetti | 6 | 66.6 mg/kg | 1, 7, 30 dpi | 1dpi = 29<br>7dpi = 68<br>30dpi = 30 | 1,2, S1-2 |
| Nestin::CreER<sup>T2</sup> | MADM | 6 | 233.1 mg/kg | 30, 60 dpi | 30dpi = 34<br>60dpi = 53 | 1,2, S1-2 |
| Nestin::CreER<sup>T2</sup> | Confetti | 12 | 99.6 mg/kg | 30, 60 dpi | 30dpi = 68<br>60dpi = 45 | 1,2, S1-2 |

**Table S2.**
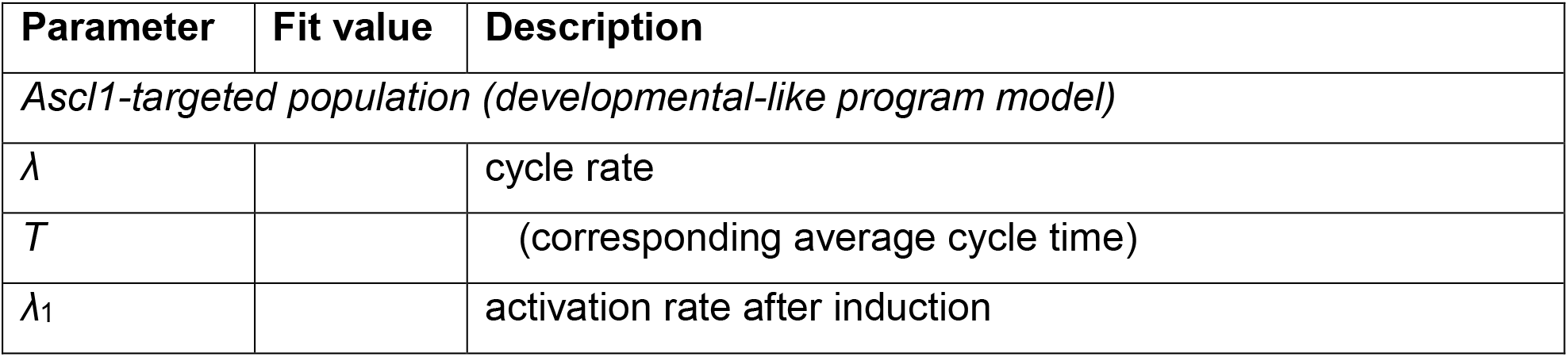
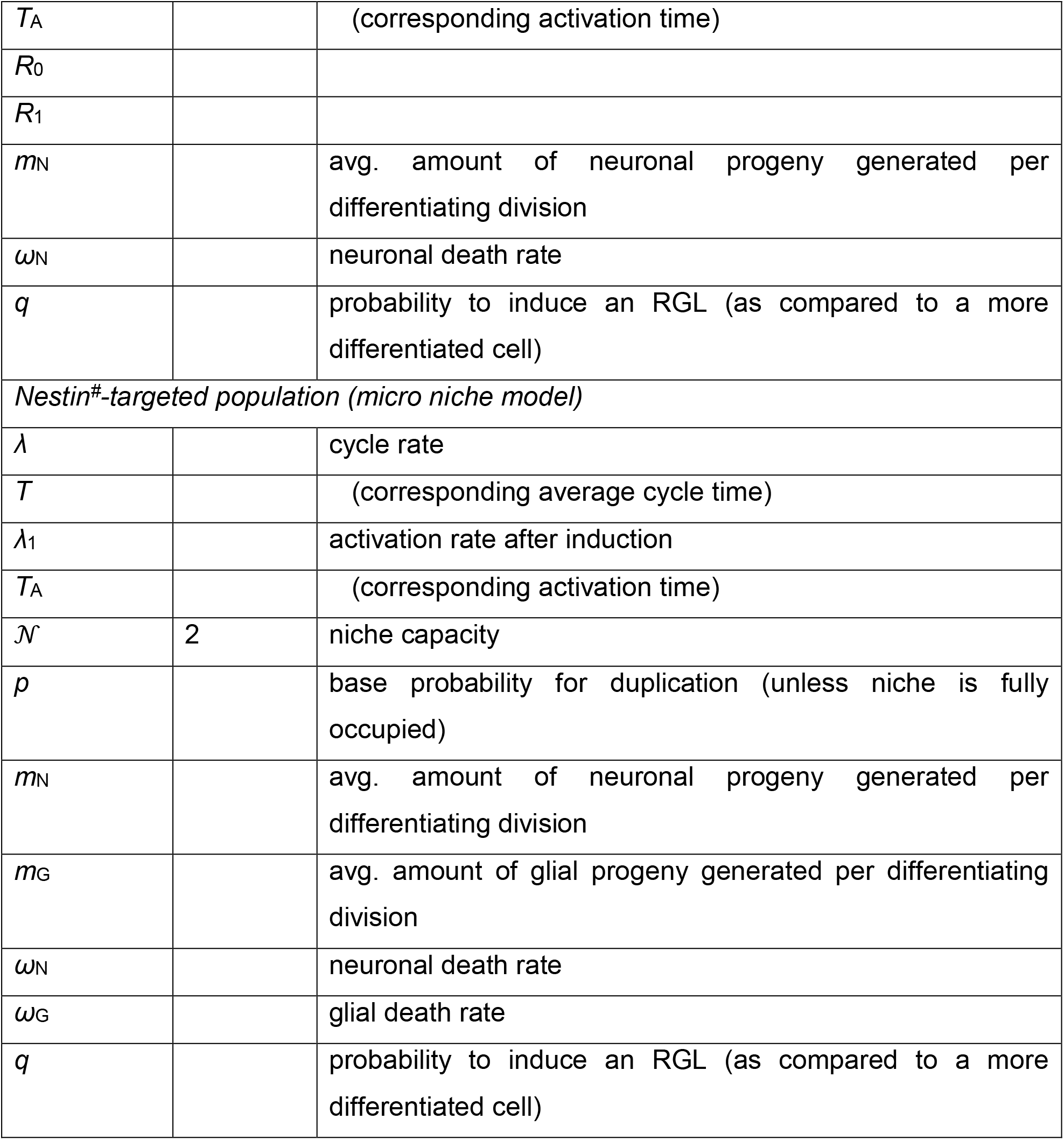
Parameter list and best fit parameters for the theoretical mode

| Parameter | Fit value | Description |
| --- | --- | --- |
| *Ascl1-targeted population (developmental-like program model)* |  |  |
| $\lambda$ | | cycle rate |
| $T$ | | (corresponding average cycle time) |
| $\lambda_1$ | | activation rate after induction |
| $T_A$ | | (corresponding activation time) |
| $R_0$ | | |
| $R_1$ | | |
| $m_N$ | | avg. amount of neuronal progeny generated per differentiating division |
| $\omega_N$ | | neuronal death rate |
| $q$ | | probability to induce an RGL (as compared to a more differentiated cell) |
| *Nestin<sup>#</sup>-targeted population (micro niche model)* |  |  |
| $\lambda$ | | cycle rate |
| $T$ | | (corresponding average cycle time) |
| $\lambda_1$ | | activation rate after induction |
| $T_A$ | | (corresponding activation time) |
| $\mathcal{N}$ | 2 | niche capacity |
| $p$ | | base probability for duplication (unless niche is fully occupied) |
| $m_N$ | | avg. amount of neuronal progeny generated per differentiating division |
| $m_G$ | | avg. amount of glial progeny generated per differentiating division |
| $\omega_N$ | | neuronal death rate |
| $\omega_G$ | | glial death rate |
| $q$ | | probability to induce an RGL (as compared to a more differentiated cell) |

**Table S3.**
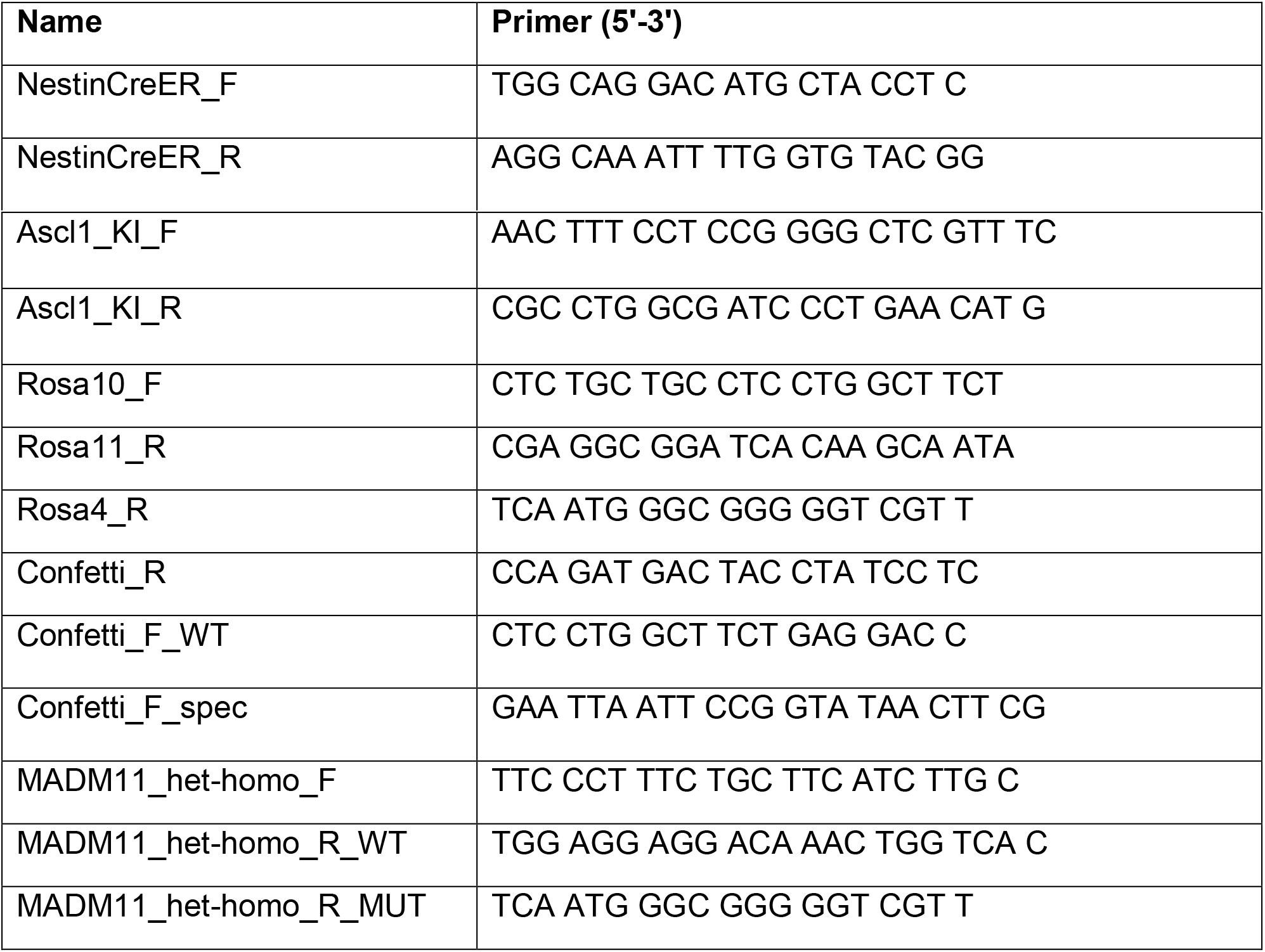
Primer sets that were used for genotyping

**Table S4.** Number of clones for clonal analysis. Summary of number of all clones, NSC-containing clones, animals across different ages and tracing time points for clonal analysis for Figures 1–3 and Supplementary Figures S1–S3.

|  | **Total number of clones** | **Total number of NSC-containing clones** | **Total number of non NSC-containing clones** | **Number of animals** |
| --- | --- | --- | --- | --- |
| **Nestin<sup>#</sup>-2mo** |  |  |  |  |
| 2 dpi | 47 | 36 | 11 | 11 |
| 7 dpi | 40 | 34 | 6 | 3 |
| 30 dpi | 54 | 45 | 9 | 6 |
| 60 dpi | 69 | 47 | 22 | 11 |
| 120 dpi | 69 | 28 | 41 | 5 |
| 365 dpi | 30 | 10 | 20 | 4 |
| **Nestin<sup>#</sup>-6mo** |  |  |  |  |
| 1 dpi | 29 | 19 | 10 | 3 |
| 7 dpi | 68 | 44 | 24 | 4 |
| 30 dpi | 64 | 53 | 11 | 5 |
| 60 dpi | 53 | 35 | 19 | 6 |
| **Nestin<sup>#</sup>-12mo** |  |  |  |  |
| 30 dpi | 68 | 46 | 22 | 3 |
| 60 dpi | 46 | 38 | 8 | 3 |
|  |  |  | Clones | Animals |
| Total of **Nestin<sup>#</sup>-lineage tracing** |  |  | 637 | 64 |
|  | **Total**<br>**number of**<br>**clones** | **Total number of**<br>**NSC-containing**<br>**clones** | **Total number**<br>**of non NSC-**<br>**containing**<br>**clones** | **Number of**<br>**animals** |
| **Ascl1<sup>#</sup>-2mo** |  |  |  |  |
| 1dpi | 85 | 71 | 14 | 7 |
| 3 dpi | 83 | 61 | 22 | 6 |
| 7 dpi | 93 | 65 | 28 | 6 |
| 30 dpi | 70 | 39 | 31 | 4 |
| 60 dpi | 59 | 24 | 35 | 4 |
| 120 dpi | 80 | 23 | 57 | 7 |
| **Ascl1<sup>#</sup>-6mo** |  |  |  |  |
| 1dpi | 30 | 15 | 15 | 3 |
| 3dpi | 89 | 30 | 59 | 5 |
| 7dpi | 101 | 57 | 44 | 9 |
| 30dpi | 78 | 20 | 58 | 6 |
| **Ascl1<sup>#</sup>-12mo** |  |  |  |  |
| 7dpi | 52 | 28 | 24 | 3 |
| 30dpi | 104 | 34 | 70 | 6 |
|  |  |  | Clones | Animals |
| Total of **Ascl1<sup>#</sup>-lineage tracing** |  |  | 924 | 66 |
| *Total number of clones and animals* |  |  | **1561** | **130** |

**Table S5. (Separate file)**

Quality metrics of RNA-seq libraries

**Table S6. (Separate file)**

Gene expression matrices

**Table S7. (Separate file)**

Differential gene expression of qNSC between young and old

**Table S8. (Separate file)**

GO term enrichment and gene presence matrix

**Table S9. (Separate file)**

Differential gene expression of comparative stem cell niches between young and aged

**Table S10. (Separate file)**

Upregulated GO term enrichment across stem cell niches and gene presence matrix

**Table S11. (Separate file)**

Downregulated GO term enrichment across stem cell niches and gene presence matrix

**Table S12. (Separate file)**

GO term co-enrichment tables

Codes for computational modeling and scRNA-seq analysis are available upon request.

**Figure S1.**
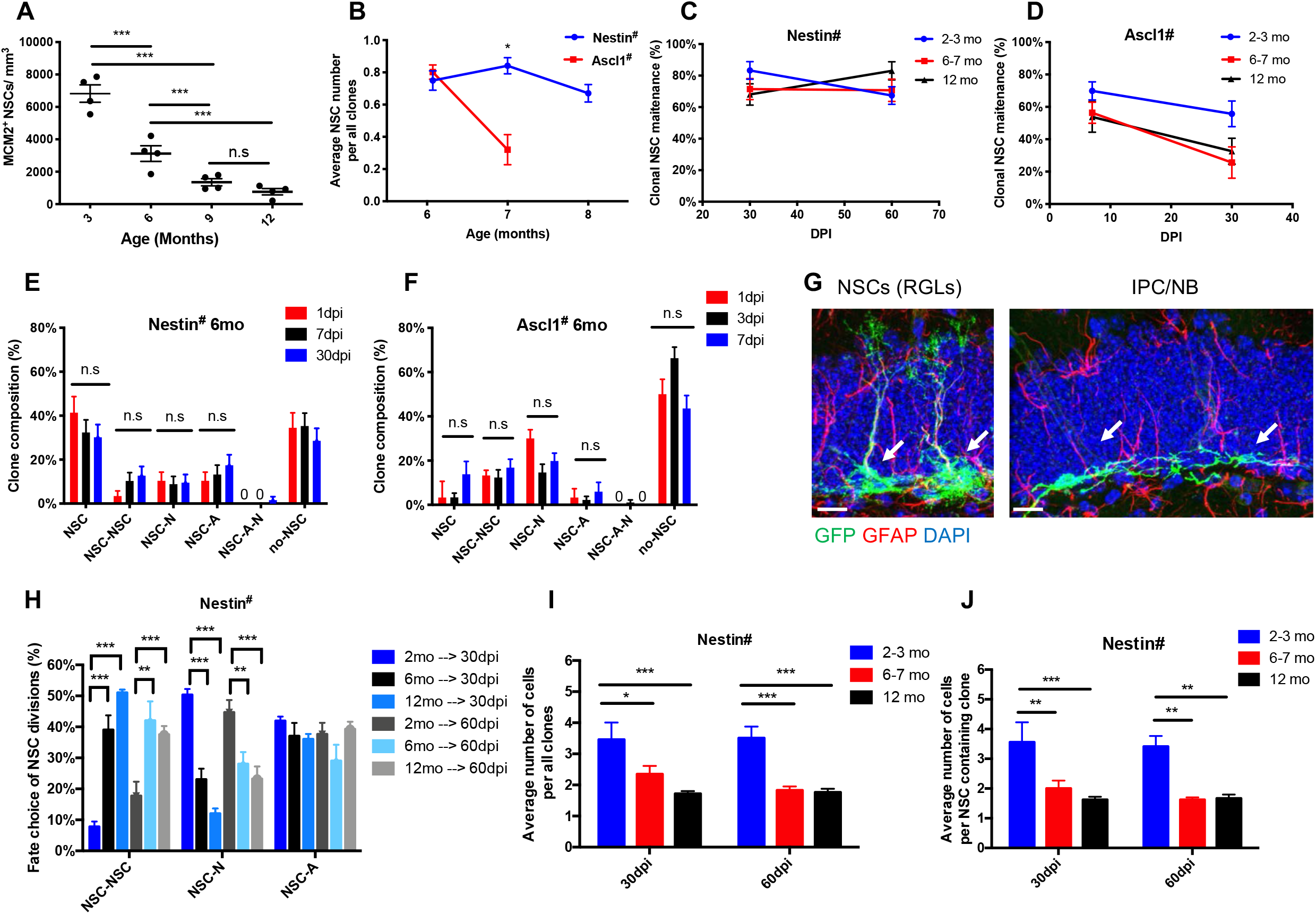
Population analysis and clonal lineage-tracing of individual NSCs in the adult mouse dentate gyrus, related to Figure 1 and 2. (A) Quantification of the total number of active NSCs in the dentate gyrus across ages. N=4-6 mice; Values represent mean ± SEM. *p<0.05, **p<0.01, ***p<0.001, n.s. - not significant; ANOVA with Bonferroni post-hoc test. (B) Quantification of NSC homeostasis duration at 6-month-old. Ascl1#-NSCs are rapidly depleted, while Nestin#-NSCs maintain as stem cells for months before eventually differentiating. Values represent mean ± SEM. (N = 29-104 clones, see details in Table S4). *p<0.05, **p<0.01, ***p<0.001, n.s. - not significant; two-way ANOVA with Tukey’s multiple comparisons test. (C) Clonal NSC maintenance from Nestin# clones acquired across multiple ages (2, 6, 12 months old animals) at multiple days post tamoxifen injection (detailed information Table S4). Shown is a summary quantification of the percent of clones which contains NSCs. (D) Clonal NSC maintenance from Ascl1# clones acquired across multiple ages (2, 6, 12 months old animals) at multiple days post tamoxifen injection (detailed information Table S4). Shown is a summary quantification of the percent of clones which contains NSCs. (E) Quantification of the clone composition for Nestin# clones at day 1-, 7- and 30-days post-tamoxifen injection into 6-month-old mice. NSC = radial glia-like neural stem cells; A=Astroglial lineage; N=neuronal lineage, 0 – no observed phenotype. Values represent mean ± SEM (N = 29-64 clones, see details in Table S4). Values represent mean ± SEM (N = 29-104 clones, see details in Table S4; *p<0.05, **p<0.01, ***p<0.001, n.s. - not significant; ANOVA with Bonferroni post-hoc test); (F) Quantification of the clone composition for Ascl1# clones at day 1-, 3- and 7-days post-tamoxifen injection into 6-month-old mice. NSC = radial glia-like neural stem cells; A=Astroglial lineage; N=neuronal lineage, 0 – no observed phenotype. Values represent mean ± SEM (N = 70-85 clones, see details in Table S4). Values represent mean ± SEM (N = 29-104 clones, see details in Table S4; *p<0.05, **p<0.01, ***p<0.001, n.s. - not significant; ANOVA with Bonferroni post-hoc test); (G) Confocal images of labeled clones reveal distinct RGL and IPCs morphologies from Nestin# and Ascl1# clones acquired from 6-month-old mouse. Shown are sample confocal images of immunostaining of GFP^+^ and GFAP^+^ (for NSCs(RGLs)); GFP^+^ and GFAP^−^ (for IPC/NB). Scale bar 20μm. (H) Quantification of NSC self-renewing cell fate divisions at 30- and 60-days post-tamoxifen injection into 2, 6 and 12 months old Nestin# mice. NSC = radial glia-like neural stem cells; A=Astroglial lineage; N=neuronal lineage. Values represent mean ± SEM (N = 29-104 clones, see details in Table S4; *p<0.05, **p<0.01, ***p<0.001, n.s. - not significant; ANOVA with Bonferroni post-hoc test); (I) Quantification of average number of cells per all clones at 30- and 60-days post-tamoxifen injection into 2, 6 and 12 months old Nestin# mice. (detailed information Table S4). (N = 30-69 clones, see details Table S4; *p<0.05, **p<0.01, ***p<0.001, n.s. - not significant; ANOVA with Bonferroni post-hoc test). (J) Quantification of average number of cells per NSC-containing clones at 30- and 60-days post-tamoxifen injection into 2, 6 and 12 months old Nestin# mice. (detailed information Table S4). (N = 30-69 clones, see details Table S4; *p<0.05, **p<0.01, ***p<0.001, n.s. - not significant; ANOVA with Bonferroni post-hoc test).

**Figure S2.**
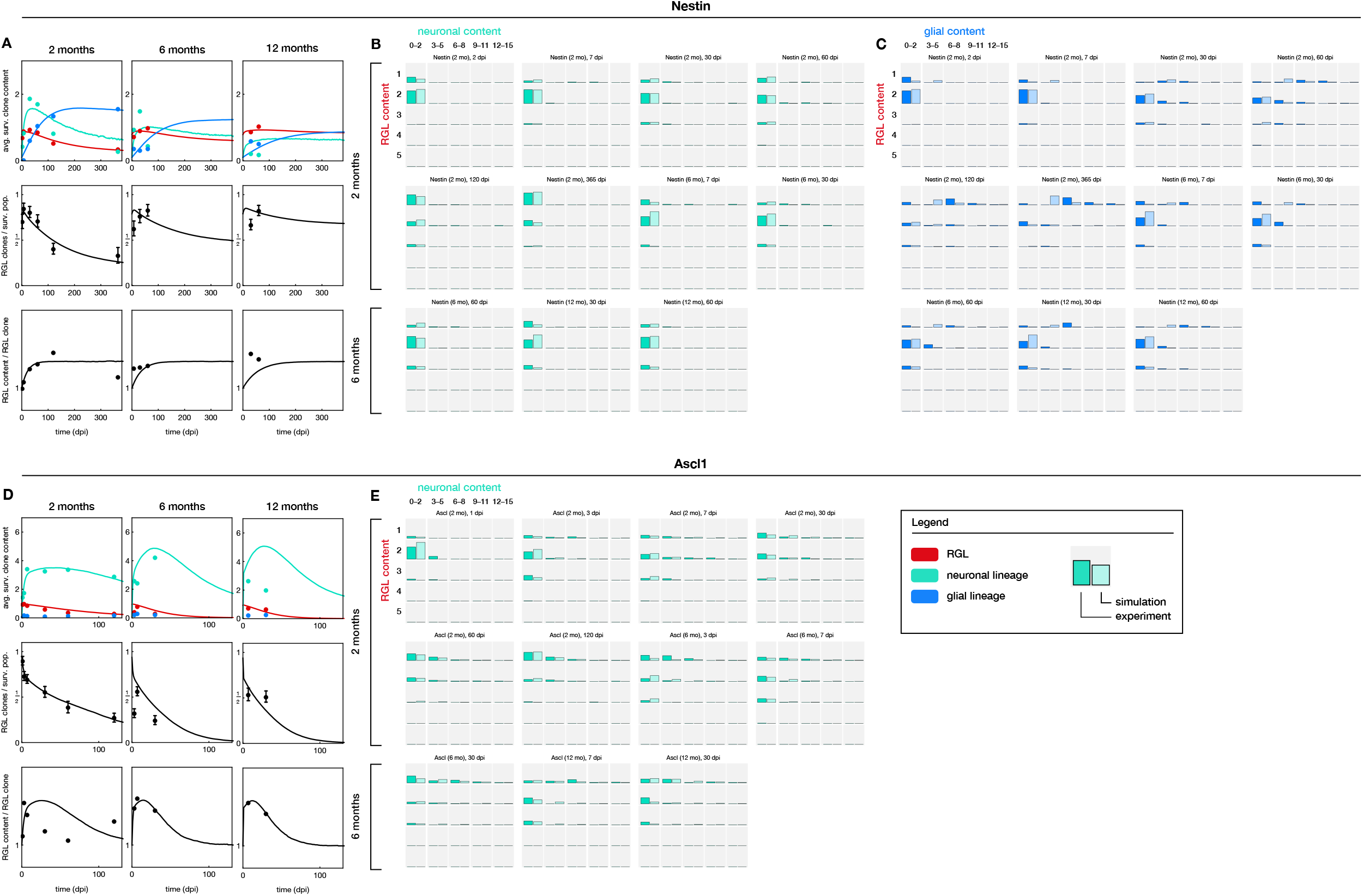
Computational modeling of Nestin#-NSC and Ascl1#-NSC, related to Figures 1 and 2. (A) Model of stochastic Nestin#-NSC fate dynamics within a small stem cell niche. NSC clonal content and survival increases in 6 and 12-month-old mice, while neuronal and astroglial content decreases compared to 2-month-old mice. Red=radial glia-like NSC content, blue=astroglial content, green=neuronal content; Line=model, dots=experimental data. (B) Least-fit squares showing correspondence between the simulated model and clonal content from Nestin#-NSCs traced in 2, 6 and 12 months-old mice. (C) Model of a developmental-like stem cell fate program in Ascl1#-NSCs. NSC clonal survival decreases in 6 and 12-month-old mice, while neuronal content increases compared to 2-month-old mice. Red=radial glia-like NSC content, blue=astroglial content, green=neuronal content. (D) Least-fit squares showing correspondence between the simulated model and clonal content from Ascl1#-NSCs traced in 2, 6 and 12 months-old mice.

**Figure S3.**
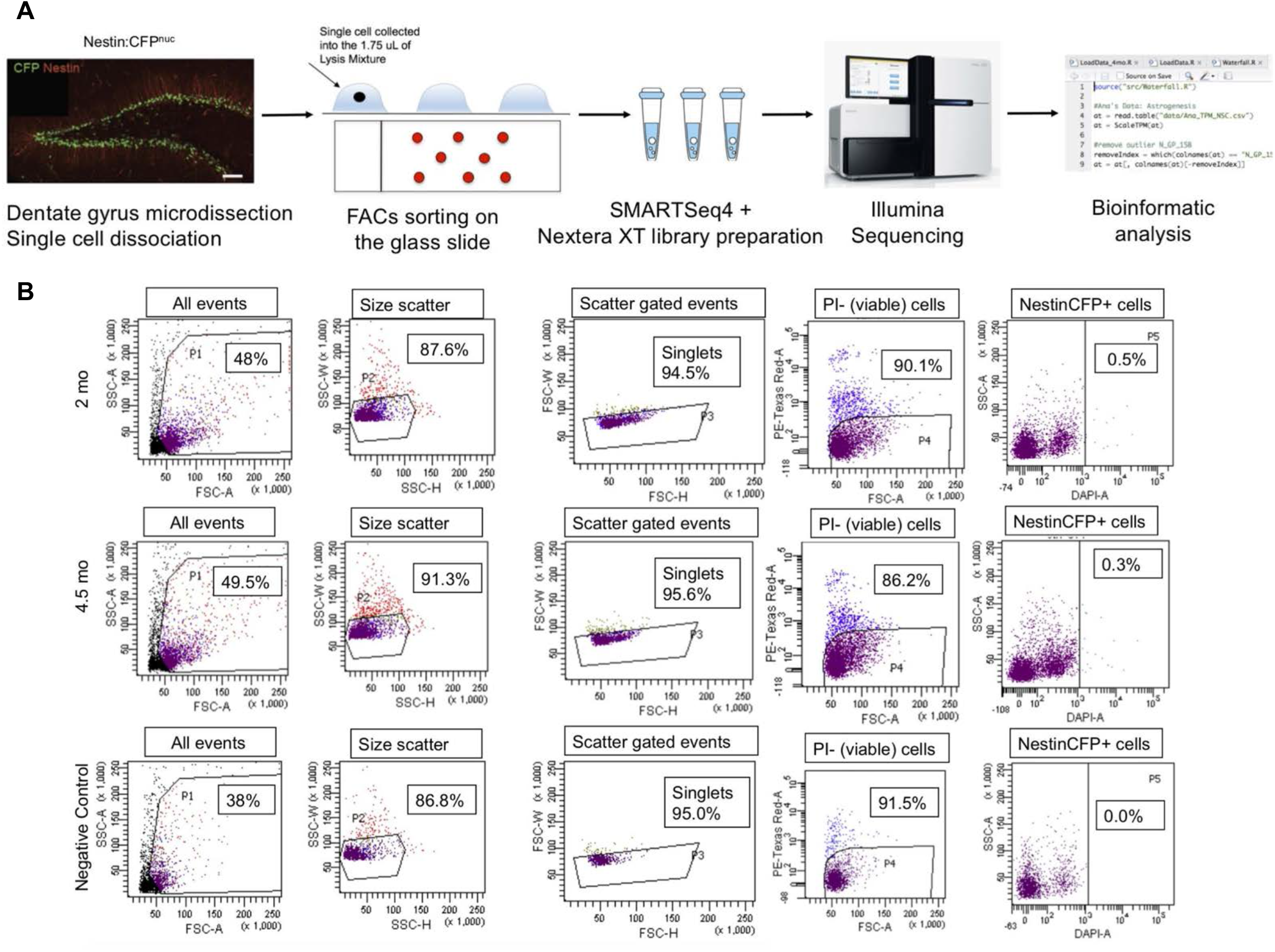
Experimental design of scRNA-seq experiment, related to Figure 3 and 4. (A) Experimental steps to generate single cell transcriptomes from 2 months-old and 4.5 months-old Nestin::CFP mice. Scale bar, 50μm. (B) Fluorescence activated cell sorting scheme used to isolate NestinCFP^+^ cells from the dentate gyrus of 2 months-old and 4.5 months-old mice. Negative controls (C57BL/6 - wild type animals) were used as indicated.

**Figure S4.**
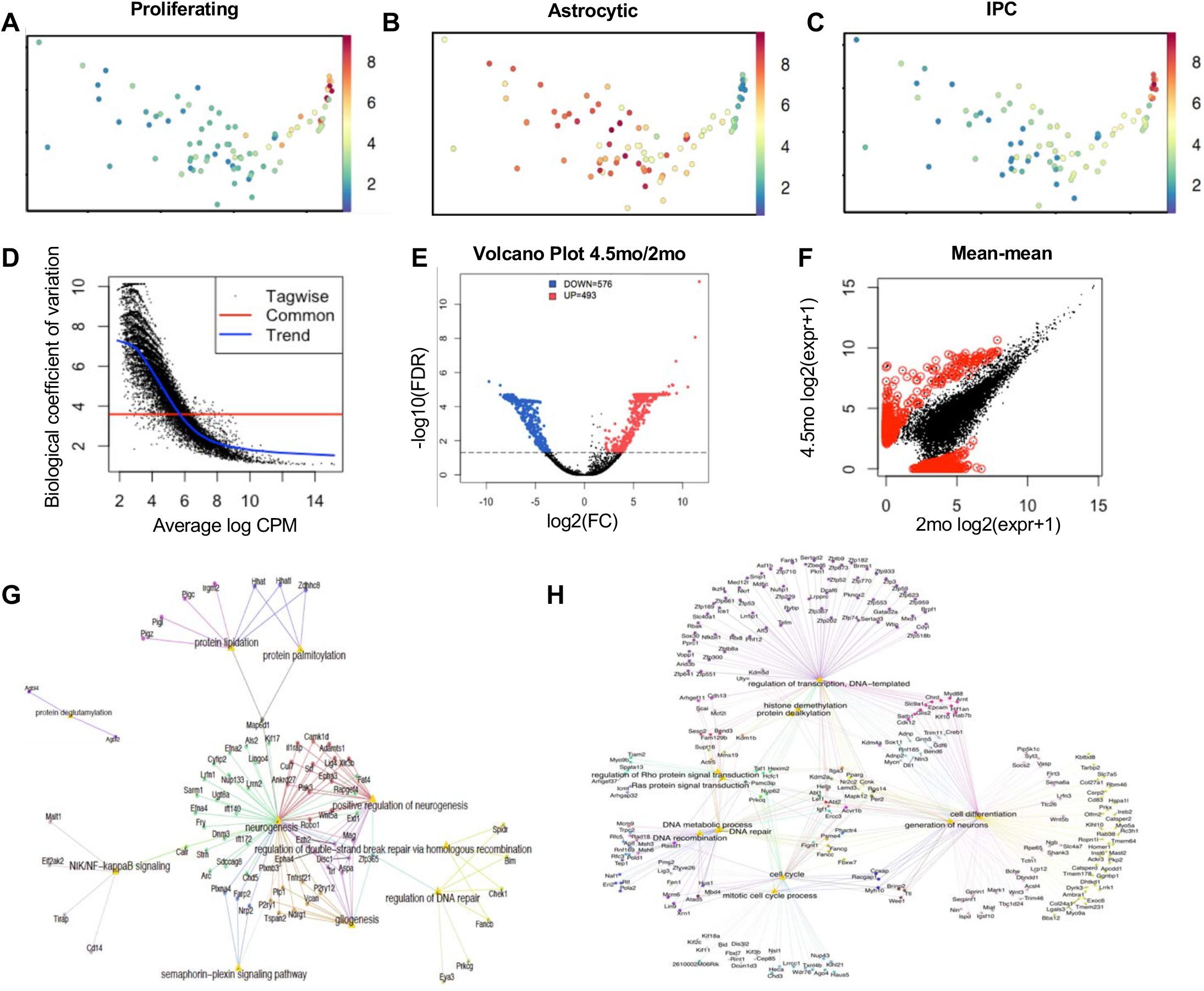
Differential gene expression analysis related to Figure 3 and 6. (A) Principle component plot highlighting proliferating cells marked by log-mean expression of PCNA, Mcm7, Cdk4, Cdk6. (B) Principle component plot highlighting astroglial properties marked by log-mean expression S100β, Aldh1l1, Megf10, Ntsr2, Acsbg1. (C) Principle component plot highlighting intermediate progenitor cells marked by log-mean expression Eomes/Tbr2, Sox11, Stmn1. (D) Biological coefficient of variation between 2 months-old and 4.5 months-old across average log counts per million (CPM). A greater degree of relative variation occurs at lower CPM levels justify a need for edgeR to call differentially expressed genes. (E) Volcano plot of edgeR-called differentially expressed genes between quiescent NSCs from 2 months-old and 4.5 months-old mice. FDR-corrected p-values are shown. (F) Mean-mean gene expression scatter plot between NSCs from 2 months-old and 4.5 months-old mice highlighting differentially expressed genes (red). (G) String network graph depicting age-related changes upregulated in 4.5-month-old vs 2-month-old quiescent NSCs. Shown genes have 1+ connections to biological processes. (H) String network graph depicting age-related changes downregulated in 4.5-month-old vs 2-month-old quiescent NSCs. Shown genes have 1+ connections to biological processes.

**Figure S5.**
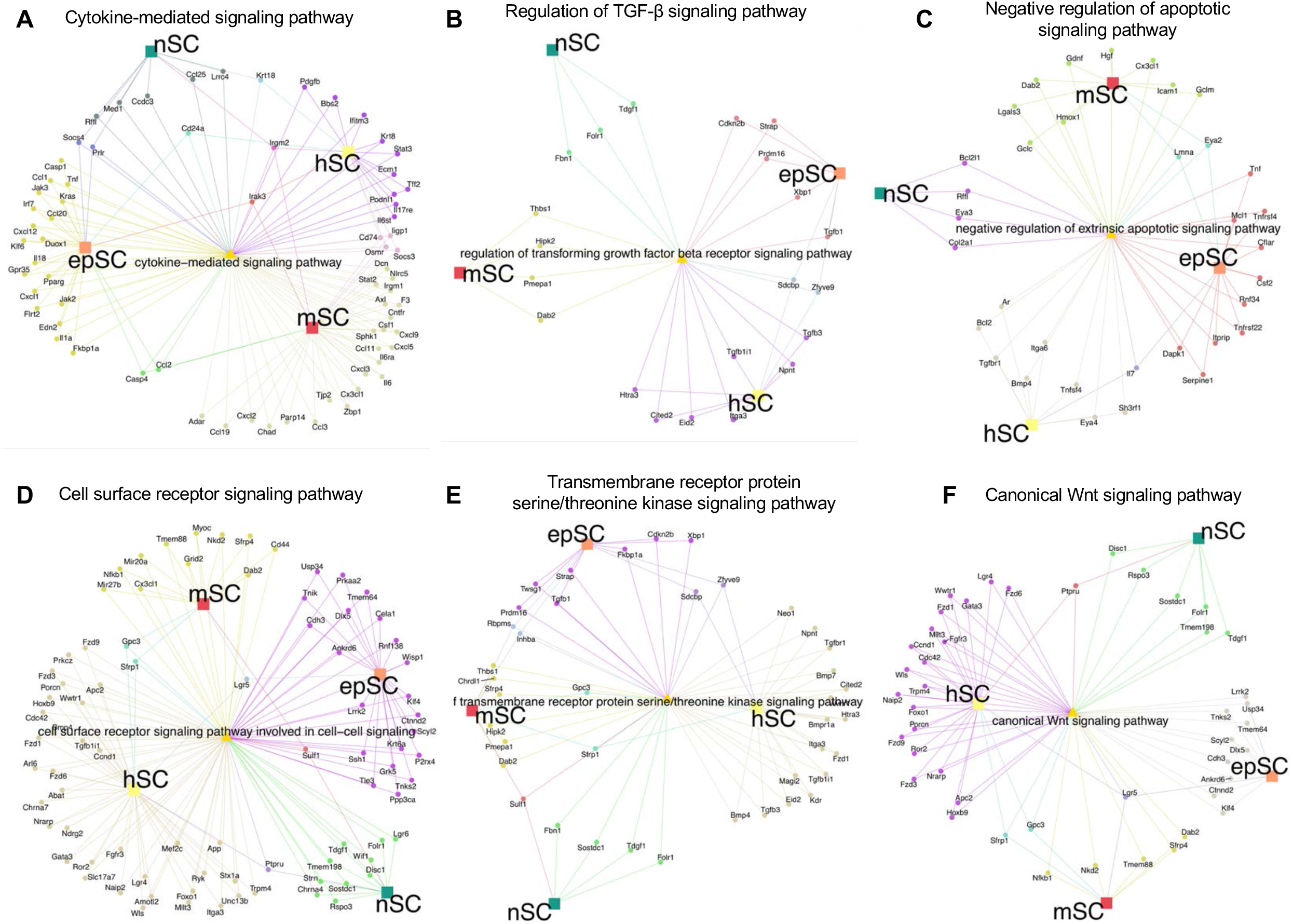
String gene network analysis for cell signaling pathways, related to Figure 4. (A) String gene network analysis showing unique or shared upregulated genes that are responsible for cytokine-mediated signaling pathway in stem cell niches. (B) String gene network analysis showing unique or shared upregulated genes that are responsible for regulation of TGFβ signaling pathway in stem cell niches. (C) String gene network analysis showing unique or shared upregulated genes that are responsible for negative regulation of apoptotic signaling pathway in stem cell niches. (D) String gene network analysis showing unique or shared upregulated genes that are responsible for cell surface receptor signaling pathway in stem cell niches. (E) String gene network analysis showing unique or shared upregulated genes that are responsible for transmembrane receptor protein serine/threonine kinase signaling pathway in stem cell niches. (F) String gene network analysis showing unique or shared upregulated genes that are responsible for canonical WNT signaling pathway in stem cell niches.

**Figure S6.**
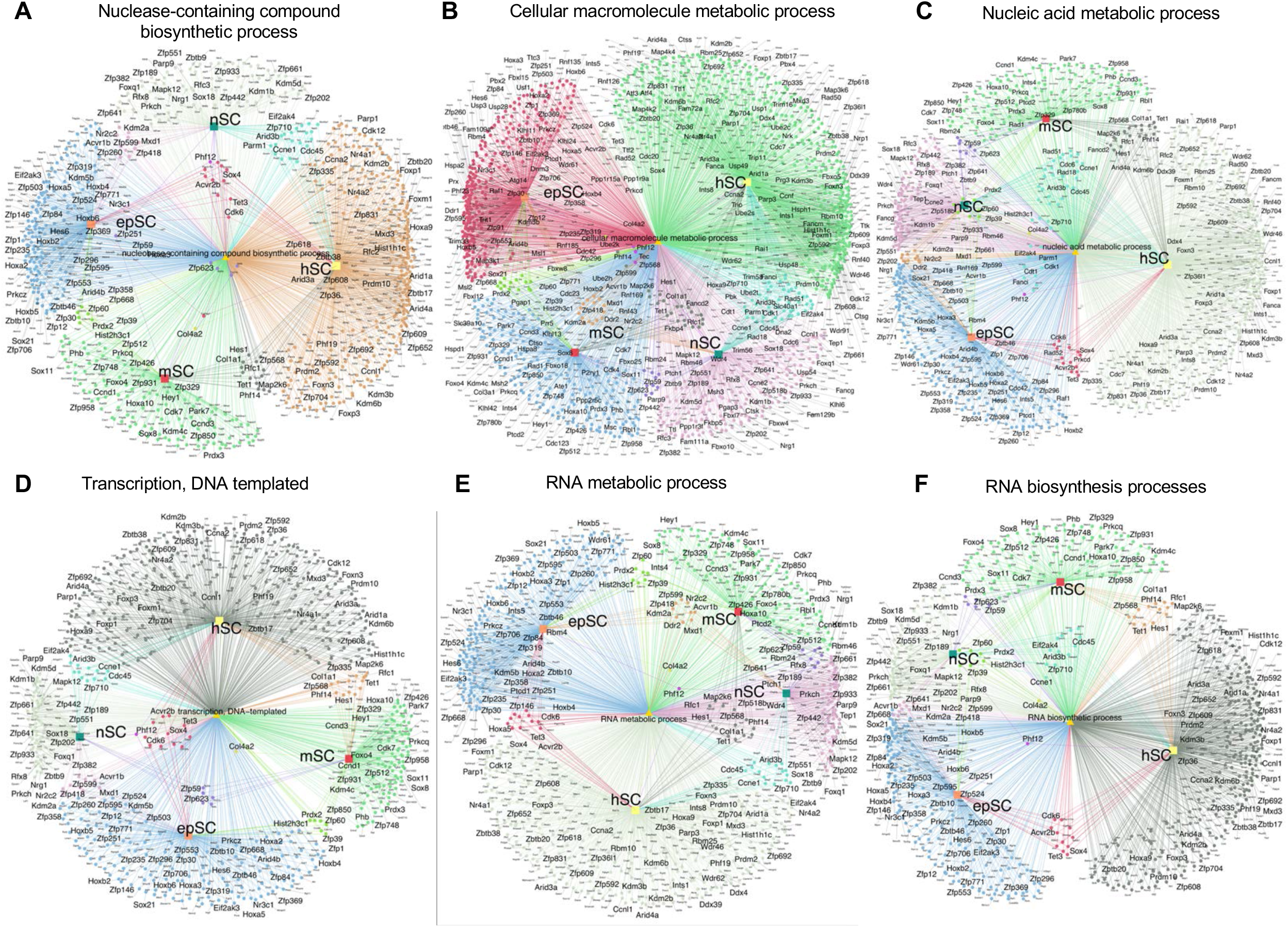
String gene network analysis for RNA homeostasis, related to Figure 4. (A) String gene network analysis showing unique or shared down-regulated genes that are responsible for nuclease – containing compound biosynthetic process in stem cell niches. (B) String gene network analysis showing unique or shared down-regulated genes that are responsible for cellular macromolecule metabolic process in stem cell niches. (C) String gene network analysis showing unique or shared down-regulated genes that are responsible for nucleic acid metabolic process in stem cell niches. (D) String gene network analysis showing unique or shared down-regulated genes that are responsible for transcription, DNA-templated in stem cell niches. (E) String gene network analysis showing unique or shared down-regulated genes that are responsible for RNA metabolic process in stem cell niches. (F) String gene network analysis showing unique or shared down-regulated genes that are responsible for RNA biosynthetic processes in stem cell niches.

**Figure S7.**
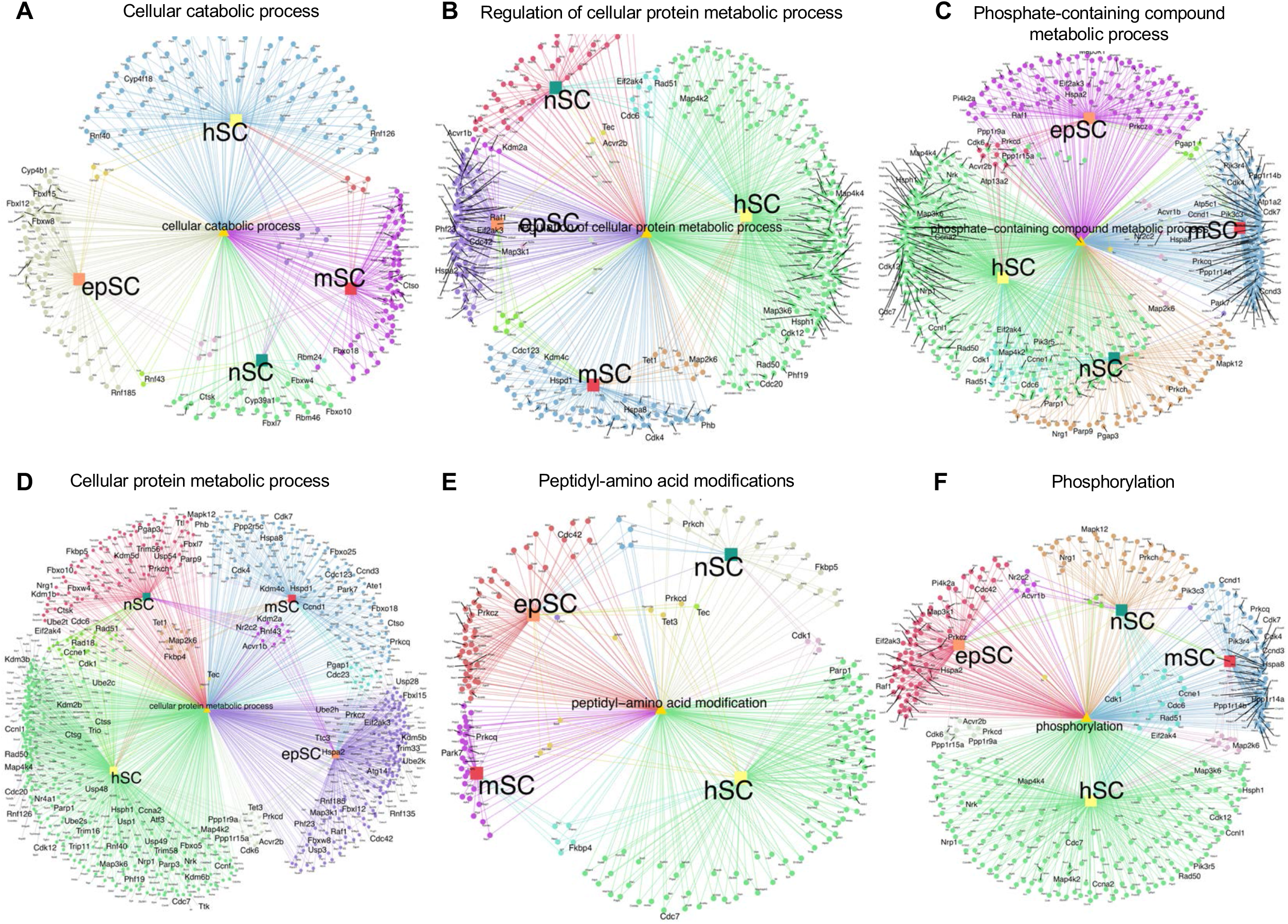
String gene network analysis for protein homeostasis, related to Figure 4. (A) String gene network analysis showing unique or shared down-regulated genes that are responsible for cellular catabolic process in stem cell niches. (B) String gene network analysis showing unique or shared down-regulated genes that are responsible for regulation of cellular protein metabolic process in stem cell niches. (C) String gene network analysis showing unique or shared down-regulated genes that are responsible for phosphate – containing compound metabolic process in stem cell niches. (D) String gene network analysis showing unique or shared down-regulated genes that are responsible for cellular protein metabolic process in stem cell niches. (E) String gene network analysis showing unique or shared down-regulated genes that are responsible for peptidyl-amino acid modifications in stem cell niches. (F) String gene network analysis showing unique or shared down-regulated genes that are responsible for phosphorylation in stem cell niches.

